# The insulin receptor adaptor IRS2 is an APC/C substrate that promotes cell cycle protein expression and a robust spindle assembly checkpoint

**DOI:** 10.1101/829572

**Authors:** Sandhya Manohar, Qing Yu, Steven P. Gygi, Randall W. King

## Abstract

Insulin receptor substrate 2 (IRS2) is an essential adaptor that mediates signaling downstream of the insulin receptor and other receptor tyrosine kinases. Transduction through IRS2-dependent pathways is important for coordinating metabolic homeostasis, and dysregulation of IRS2 causes systemic insulin signaling defects. Despite the importance of maintaining proper IRS2 abundance, little is known about what factors mediate its protein stability. We conducted an unbiased proteomic screen to uncover novel substrates of the Anaphase Promoting Complex/Cyclosome (APC/C), a ubiquitin ligase that controls the abundance of key cell cycle regulators. We found that IRS2 levels are regulated by APC/C activity and that IRS2 is a direct APC/C target in G_1_. Consistent with the APC/C’s role in degrading cell cycle regulators, quantitative proteomic analysis of IRS2-null cells revealed a deficiency in proteins involved in cell cycle progression. We further show that cells lacking IRS2 display a weakened spindle assembly checkpoint in cells treated with microtubule inhibitors. Together, these findings reveal a new pathway for IRS2 turnover and indicate that IRS2 is a component of the cell cycle control system in addition to acting as an essential metabolic regulator.

## Introduction

The insulin and insulin-like growth factor 1 receptors (IR/IGF1R) are receptor tyrosine kinases that control metabolism, differentiation, and growth. Upon ligand binding at the cell surface, the activated IR/IGF1R undergoes a conformational change that allows it to auto-phosphorylate tyrosine residues on its cytoplasmic subunits [1]. This facilitates the recruitment and phosphorylation of insulin receptor substrate (IRS) proteins, which serve as scaffolds to initiate downstream signaling [2]. Two major pathways that are stimulated by this cascade are the PI3K-AKT and Ras-Raf-MAPK pathways, which coordinate metabolic homeostasis and growth, among other functions [1].

The most physiologically important and ubiquitously expressed IRS proteins are IRS1 and IRS2. Though IRS1 and IRS2 share similar structural and functional features, they have complementary roles and expression patterns that depend on tissue type and physiological state [1]. These differences are illustrated by divergent phenotypes in knockout mice: whereas IRS1 knockout mice exhibit insulin resistance that is compensated by increased pancreatic β cell mass, IRS2 knockout mice exhibit β cell failure and resultant diabetes [3]. Distinct roles for IRS1 and IRS2 can also be observed within the same tissue. For example, in skeletal muscle, IRS1 is required for glucose uptake and metabolism, whereas IRS2 is important for lipid uptake and metabolism [4, 5]. Furthermore, recent work has shown that the ratio of IRS1 to IRS2 is important for hepatic glucose metabolism [6]. Thus, maintaining proper IRS1 and IRS2 levels is critical for systemic and cellular homeostasis.

The ubiquitin-mediated proteolysis of IRS proteins is important for restraining signaling through the IR/IGF1R. For example, both IRS proteins are targeted for proteasomal destruction following persistent insulin or IGF1 stimulation in a negative feedback loop that attenuates PI3K-AKT signaling [2, 7]. In mice, removal of a ubiquitin ligase that is responsible for IGF1-induced degradation of IRS1 enhances insulin sensitivity and increases plasma glucose clearance [7]. Though several ubiquitin ligases have been reported to control IRS1’s proteasome-dependent degradation [8-12], only SOCS1/3 have been implicated in driving IRS2 turnover [11]. This is an intriguing disparity because hepatic IRS1 remains stable between fasting and feeding whereas IRS2 levels drop after feeding [13], suggesting that IRS2 is less stable than IRS1 in some physiological contexts. Because SOCS1/3 also targets IRS1, there are no reports of ubiquitin ligases that target IRS2 but not IRS1, leaving a gap in our knowledge of how IRS1 and IRS2 are differentially regulated by the ubiquitin proteasome system.

The Anaphase-Promoting Complex/Cyclosome (APC/C) is a 1.2 mDa ubiquitin ligase that targets key cell cycle related proteins for destruction by the proteasome [14, 15]. To transfer ubiquitin to its substrates, the APC/C works with one of two co-activators: Cdc20 during M-phase or Cdh1 during G_1_. These co-activators stimulate the catalytic activity of the APC/C and facilitate substrate recognition. APC/C^Cdc20^ and APC/C^Cdh1^ recognize substrates via short degron motifs in unstructured protein regions called destruction boxes (D-boxes) and KEN-boxes. An additional degron, called the ABBA motif, is used only used by APC/C^Cdc20^ in metazoan cells [14-16].

To probe the substrate landscape of the APC/C, we conducted an unbiased proteomic screen by acutely blocking APC/C^Cdh1^ activity with small molecule APC/C inhibitors (apcin and proTAME) [17, 18] in G_1_ cells. Using this approach, we uncovered diverse putative APC/C^Cdh1^ substrates, including IRS2. We demonstrate that IRS2, but not IRS1, is a direct target of APC/C^Cdh1^, thereby establishing a novel mode by which IRS1 and IRS2 are differentially regulated. Using IRS2 knockout cell lines, we show that IRS2 is important for the expression of proteins involved in cell cycle progression. We further show that genetic deletion of IRS2 perturbs spindle assembly checkpoint function. Taken together, these data establish a role for IRS2 in normal cell cycle progression, revealing new connections between an essential component of the growth factor signaling network and cell cycle regulation.

## Results

### Chemical proteomics reveals proteins whose abundances are APC/C^Cdh1^ regulated

To identify novel substrates and pathways regulated by APC/C^Cdh1^, we designed an experiment that coupled small molecule inhibition of the APC/C in G_1_ cells to high resolution tandem mass tag (TMT)-based quantitative proteomics (**Figure 1A**). Blocking Cdk4/6 activity inhibits Rb phosphorylation, causing cells to arrest at the G_1_ restriction point [19]. Thus, to generate a homogeneous population of G_1_ cells, we treated asynchronous hTERT-RPE1 cells bearing fluorescent ubiquitination-based cell cycle indicator (FUCCI) constructs [20] with the Cdk4/6 inhibitor palbociclib. Following G_1_ arrest, cells were acutely treated with a combination of APC/C inhibitors (6 µM proTAME + 50 µM apcin) or vehicle (DMSO) for 8 hours. Cells were then collected for proteomic analysis with the expectation that APC/C-regulated proteins would be stabilized in cells treated with APC/C inhibitors compared to control cells (**Figure 1A**). The combined use of proTAME and apcin results in robust inhibition of the APC/C [17], which guided our decision to use this treatment scheme. Moreover, this scheme was designed to specifically identify APC/C^Cdh1^ substrates rather than APC/C^Cdc20^ substrates since APC/CCdh1 degrades Cdc20 during G_1_ phase [21]. Illustrating this point, Cdc20 expression was strongly reduced in G_1_ palbociclib-arrested cells (**Figure 1B**). The expression of cyclins A and B was also reduced, consistent with a G_1_ block.

**Figure 1:**
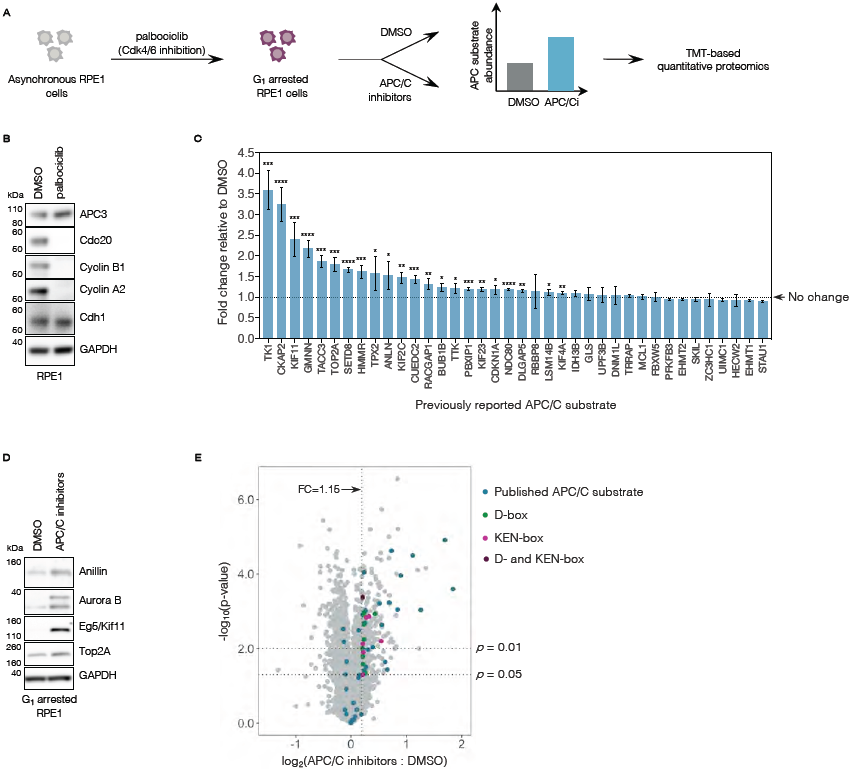
High resolution chemical proteomics reveals proteins whose abundances are APC/C regulated. (A) Workflow for the chemical proteomics experiment described in this study. Asynchronous RPE1 cells were arrested in 1 µM palbociclib (a Cdk4/6 inhibitor) for 20 hours, at which point they were acutely treated with either DMSO or a combination of 6 µM proTAME + 50 µM apcin (referred to as “APC/C inhibitors” or “APC/Ci”). Cells were then collected at time 0 (the time of drug addition) or 8 hours after drug addition and were harvested for TMT-based proteomic identification and quantification. Samples were analyzed in biological triplicate within a 10-plex TMT label set, with the 10^th^ channel used as a bridge. (B) Asynchronous RPE1 cells were treated with either DMSO or 1 µM palbociclib for 20 hours. Cells were harvested, and lysates were analyzed by immunoblot for the indicated proteins. (C)Previously reported APC/C substrates that were identified in this study are plotted with their observed fold change in the APC/C inhibitor treated sample (APC/Ci) relative to the DMSO treated sample. Error bars represent the standard deviation (SD) between the three biological replicates measured by MS. Asterisks indicate an abundance increase over control that is statistically significant (* : *p* < 0.05 ; ** : *p* < 0.01 ; *** : *p* < 0.001 ; **** : *p* < 0.0001) (D) Previously reported APC/C substrates that were identified as increasing by mass spectrometry in G_1_ RPE1 cells treated with APC/C inhibitors were validated by immunoblot for selected proteins. (E) Volcano plot highlighting all published APC/C substrates identified in this study (blue) as well as proteins that (1) contain a high probability D- and/or KEN-box (D-box = green, KEN-box = pink, D- and KEN-boxes = purple), (2) increase ≥1.15-fold under APC inhibition, (3) were identified by >1 peptide, and (4) have a *p*-value < 0.05.

The experimental approach outlined in **Figure 1A** was validated using the FUCCI reporter system. This system relies on the expression of two stably integrated fluorescent fusion proteins—mAG1-geminin (1-110) and mCherry-Cdt1 (30-120)—to monitor the activity of endogenous cell cycle-related ubiquitin ligases APC/C^Cdh1^ and SCF^Skp2^, respectively [20]. As expected, cells treated with palbociclib showed a reduction in mAG1-geminin (1-110) fluorescence over time due to APC/C^Cdh1^ activity while mCherry-Cdt1 (30-120) intensity was increased, indicating G_1_ arrest (**Figures S1A-S1B**). The addition of APC/C inhibitors in palbociclib-arrested cells rescued mAG1-geminin (1-110) levels (**Figures S1C-S1D**), confirming that this workflow stabilizes APC/C targets. Notably, cells released from palbociclib-mediated arrest accumulate mAG1-geminin (1-110) more rapidly than palbociclib-arrested cells treated with APC/C inhibitors (**Figure S1E**), indicating that APC/C inhibition is likely insufficient to trigger cell cycle re-entry under these conditions.

Using TMT-coupled quantitative proteomics, we identified and quantified relative abundances for ∼8000 human proteins in G_1_-arrested cells treated with or without APC/C inhibitors in biological triplicate (**Supplementary Table S1**). Notably, we detected 38 previously reported APC/C substrates in our dataset (**Figures 1C-1E; Supplementary Table S2)**. Of these, 22 increased significantly (*p* < 0.05) under conditions of APC/C inhibition. We validated these findings in the context of several previously reported substrates by immunoblot (**Figure 1D**). As an internal control, we detected a significant increase (*p* = 3.2 × 10^−5^) in the abundance of peptides derived from the N-terminal 110 amino acids of geminin (GMNN). These residues are shared with the mAG1-geminin (1-110) reporter expressed in this cell line, confirming earlier fluorescence-based validation of our experimental system.

While the majority of the previously reported APC/C^Cdh1^ substrates that were quantified in our G_1_ proteomic experiment were stabilized following APC/C inhibition, some remained constant. There are several possible explanations for this result. First, for proteins that were identified based on few peptides, inadequate quantification may have resulted in inaccurate abundance assignments. Second, some substrates may be APC/C^Cdh1^-accessible only under conditions or in tissue types that were not met by the experimental parameters that we used. Third, some proteins (e.g. FBXW5, ZC3HC1) [22, 23] were proposed to be APC/C^Cdh1^ substrates based on results obtained in Cdh1 overexpression systems, indicating that APC/C^Cdh1^ activity may be sufficient but not necessary to control their levels.

Of the 38 previously reported APC/C substrates that we identified, the median fold change under APC/C inhibition compared to DMSO was 1.15. Based on this, to identify new APC/C substrates, we screened for proteins that: (1) had a fold change ≥1.15 under APC/C inhibition, (2) were identified and quantified based on >1 peptide, and (3) had a *p*-value < 0.05 across the three biological replicates measured in this experiment. This narrowed our analysis to a subset of 204 proteins (**Supplementary Table S3**). Because the APC/C recognizes substrates based on D-box motifs (RxxL or the extended motif RxxLxxxxN) and KEN-box motifs (KEN), we used the SLiMSearch (Short Linear Motif Search) degron prediction tool [16, 24] to scan this 204-protein subset for proteins that contain these sequences. In order to classify a putative D- or KEN-box sequence as a probable physiological degron, we applied the following restrictions on the SLiMSearch [24] parameters: (1) similarity score ≥ 0.75; (2) consensus similarity is medium or high; (3) disorder score ≥ 0.4; (4) the putative degron must be intracellular and exist on a non-secreted protein. These cutoffs were determined based on those met by previously validated APC/C substrates (including those not identified in our dataset) and by the physical restriction that APC/C activity occurs within the cell. Based on these thresholds, our analysis identified 26 proteins as potential D- and KEN-box containing APC/C^Cdh1^substrates (**Table 1, Figure 1E**). Of these 26 proteins, 11 have previously been reported as direct APC/C substrates, validating internally that this analysis was useful for identifying APC/C substrates.

**Table 1:**
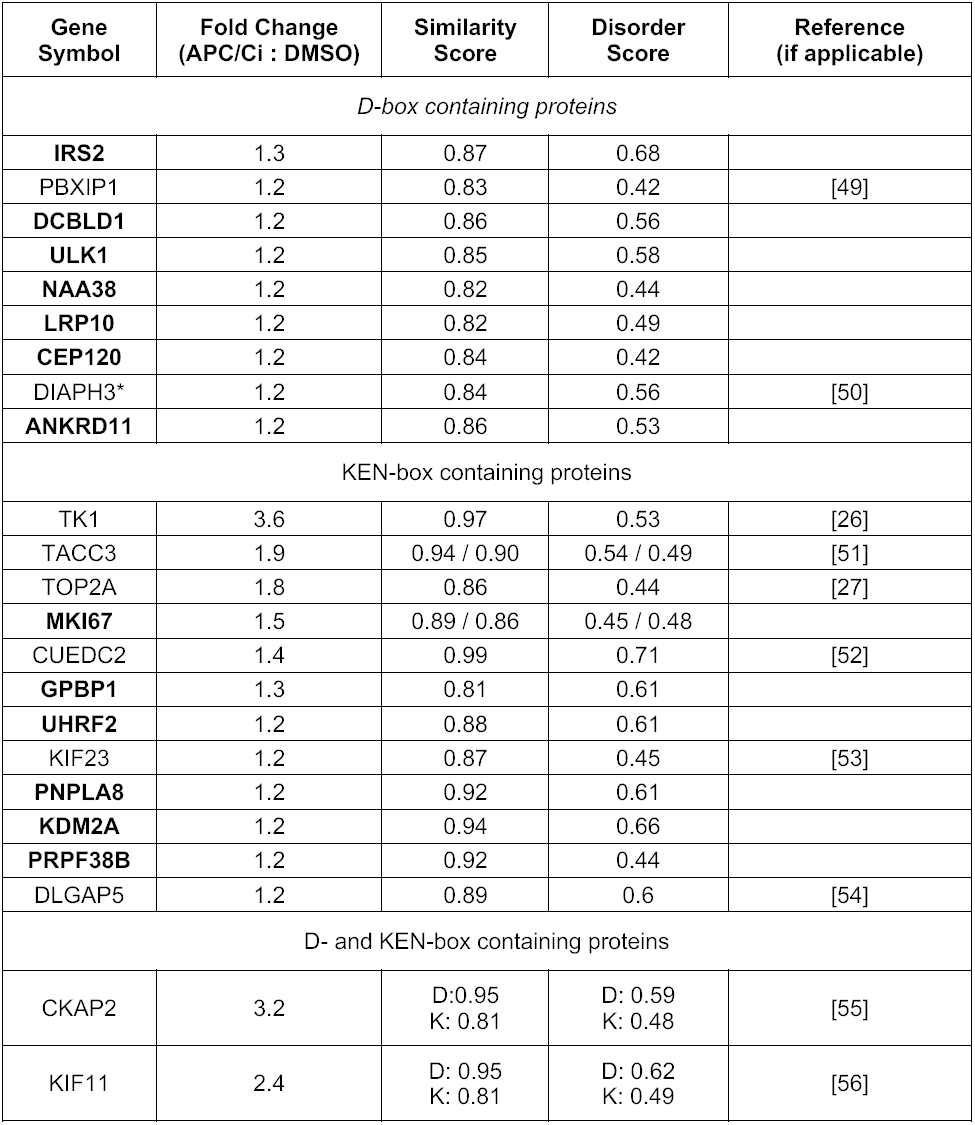
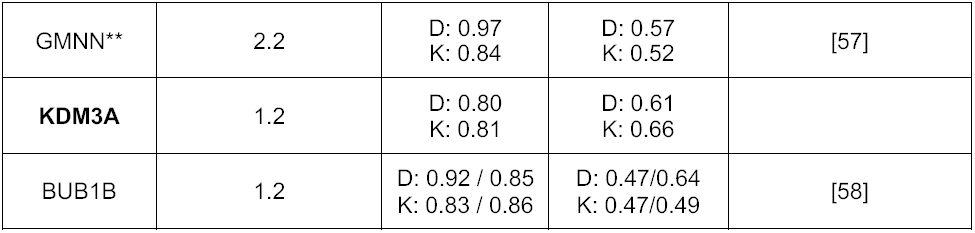
26 proteins containing high-probability D- and KEN-boxes as identified from G_1_ APC/C inhibitor proteomics. Features of the putative degron(s) found in each protein are annotated, including the SLiMSearch similarity score to other validated degrons, the similarity of the surrounding consensus sequence to other validated degrons, the disorder score for the region of the protein in which the degron is located, and the citation of the publication that reports the protein as an APC/C substrate, where applicable. *While DIAPH3/mDia2 has been shown to be ubiquitinated in a cell cycle dependent manner and was suggested as an APC/C substrate, there is no direct cell-based or biochemical evidence for this. **We cannot delineate whether the geminin peptides identified here derive from the FUCCI reporter or the endogenous protein. Proteins that have not been previously reported as APC/C substrates are shown in bold.

### IRS2 levels are controlled by Cdh1 in a proteasome-dependent manner

Examining our 26-protein putative substrate list, we focused our attention on IRS2—one of two major adaptors that promotes signaling through the insulin and insulin-like growth factor 1 receptors (IR/IGF1R). Using conditions identical to those under which the proteomics experiment was conducted, we validated that IRS2 was upregulated at the protein level under APC/C inhibition in G_1_-arrested RPE1 cells by immunoblot (**Figure 2A**). Seeking to further validate this result in a distinct physiological context, we asked whether APC/C inhibition in terminally differentiated C_2_C_12_ myotubes also increases IRS2 protein abundance. C_2_C_12_ myoblasts easily differentiate into multinucleated myotubes following serum withdrawal and supplementation with growth factors (**Figures S2A-S2B**). To validate that the APC/C is active in this system, we transfected C_2_C_12_ myoblasts with a model APC/C substrate (N-terminal fragment of cyclin B1 fused to EGFP; NT-CycB-GFP), allowed cells to differentiate into myotubes, and found that APC/C inhibition stabilized NT-CycB-GFP (**Figure S2C**). Similarly, we found that acute APC/C inhibition in myotubes also resulted in an accumulation of IRS2 protein (**Figure 2B**), thereby validating this finding from our G_1_ experiment in RPE1 cells in an independent system.

**Figure 2:**
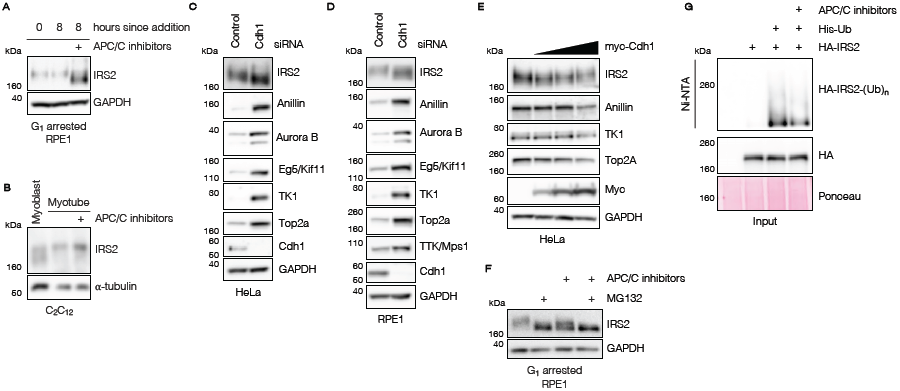
IRS2 levels are controlled by Cdh1 in a proteasome-dependent manner. (A) Cells were treated identically to what is described in **Figure 1A**, and IRS2 abundance was measured by immunoblot. (B) C_2_C_12_ myoblasts were induced to differentiate through serum withdrawal and supplementation with insulin, transferrin, and selenium (ITS). After three days of differentiation, myotubes were acutely treated with either DMSO or APC/C inhibitors (of 6 µM proTAME + 50 µM apcin). After eight hours of drug treatment, myotubes were collected and IRS2 levels from all samples were analyzed by immunoblotting. (C) – (D) Asynchronous HeLa (*C*) and RPE1 (*D*) cells were transfected with either a control or Cdh1-directed siRNA for 24 hours. Cells were allowed to grow for an additional 24 hours prior to collection and analysis of the indicated protein levels in lysate by immunoblot. (E) HeLa cells were mock transfected or transfected with increasing amounts of a plasmid encoding myc-tagged human Cdh1 for 24 hours. Cells were allowed to grow for an additional 24 hours prior to collection and analysis of the indicated protein levels by immunoblot. (F) RPE1 cells were arrested in G_1_ with 1 µM palbociclib for 20 hours. Following G_1_ arrest, cells were treated with DMSO, APC/C inhibitors (6 µM proTAME + 50 µM apcin), MG132 (10 µM), or a combination of APC/C inhibitors and MG132 for an additional 8 hours. Cells were harvested, and lysates were analyzed by immunoblot for IRS2 abundance. (G) 6x-His-tagged ubiquitin conjugates were isolated from HeLa cell lysates using Ni-NTA agarose resin. Lysates were derived from cells expressing 6x-His-ubiquitin and HA-tagged IRS2 that were treated with MG132 alone or in combination with APC/C inhibitors. Resin eluate and inputs were probed by immunoblot using an HA antibody, and Ponceau staining was used as a loading control.

To exclude the possibility that the change in IRS2 abundance that we observed following APC/C inhibition was due to off-target effects of the small molecule APC/C inhibitors, we depleted Cdh1 using RNAi to block APC/C^Cdh1^ activity in HeLa, RPE1, and asynchronous C_2_C_12_ cells (**Figure 2C-2D, Figure S2D**). We found that Cdh1 knockdown caused an accumulation of endogenous IRS2 as well as several other previously reported APC/C substrates compared to control-transfected cells (**Figure 2C-2D**).

To address whether increased APC/C^Cdh1^ activity is sufficient to reduce IRS2 levels, we overexpressed myc-tagged human Cdh1 in HeLa cells. We found that, when expressed at sufficiently high levels, Cdh1 reduced the levels of both IRS2 as well as other known APC/C^Cdh1^ substrates including anillin, TK1, and Top2a (**Figure 2E**) [25-27]. The requirement that Cdh1 be expressed at high levels to observe this effect is likely due to Cdk-dependent inhibitory phosphorylation of Cdh1 limiting its ability to activate APC/C under sub-saturating conditions [28].

We next sought to confirm that the increase in IRS2 protein observed under APC/C inhibition was due to impaired targeting of IRS2 to the proteasome. To test this, we arrested RPE1 cells in G_1_ using palbociclib and acutely treated them with APC/C inhibitors and/or a proteasome inhibitor (MG132) for 8 hours. This experiment revealed that APC/C inhibition or proteasome inhibition each resulted in an accumulation of IRS2 (**Figure 2F**). Notably, co-inhibition of the APC/C and the proteasome did not result in additional stabilization of IRS2, indicating that the increase in IRS2 we observed under APC/C inhibition was solely a consequence of its impaired proteasomal degradation. Consistent with this observation, we found that APC/C inhibition decreased the polyubiquitination of HA-tagged IRS2 in HeLa cells treated with MG132 (**Figure 2G**).

### IRS2 levels and phosphorylation fluctuate in a cell-cycle dependent manner

To test whether IRS2 levels fluctuate during the cell cycle as expected for an APC/C substrate, we synchronized HeLa cells in early S-phase by double thymidine block and tracked IRS2 protein abundance leading into mitotic entry by immunoblot (**Figure 3A**). As is typical for APC/C substrates, IRS2 levels correlated with cyclin B1 abundance and APC3 phosphorylation. To assess IRS2 levels at mitotic exit, we thymidine-nocodazole synchronized HeLa cells, released them into prometaphase, and tracked IRS2’s abundance through mitotic exit (**Figure 3B**). Again, IRS2 protein abundance correlated with cyclin B1 levels and APC3 phosphorylation. The same behavior was observed in RPE1 cells that were synchronized in late G_2_ by RO3306 treatment (Cdk1 inhibition) and tracked over the course of progression through M-phase and into G_1_ (**Figure 3C**). Based on these data, we conclude that IRS2 protein levels fluctuate in a cell cycle-dependent manner that is consistent with other known APC/C substrates. This behavior is also consistent with IRS2 being a potential APC/C^Cdc20^ substrate.

**Figure 3:**
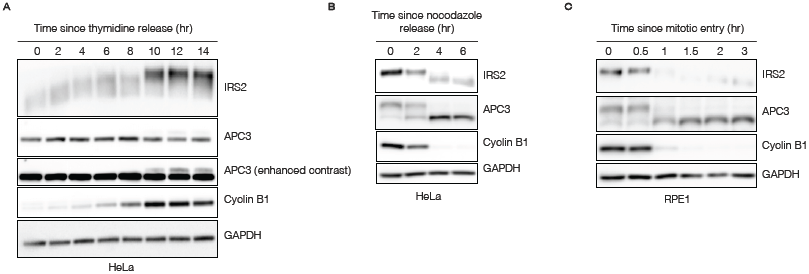
IRS2 levels and phosphorylation fluctuate in a cell-cycle dependent manner. (A) HeLa cells were synchronized by double thymidine block and released into S-phase in the presence of nocodazole. Lysates were harvested and analyzed by immunoblotting for IRS2 and cell cycle markers. (B) HeLa cells were synchronized by single thymidine-nocodazole block and released into prometaphase. Mitotic cells were collected by mitotic shake-off and re-plated. Time points were taken every two hours as cells exited M-phase. Lysates were harvested and analyzed by immunoblotting for IRS2 and cell cycle markers. (C) RPE1 cells were synchronized in G_2_ by treatment with the Cdk1 inhibitor RO3306. After 18 hours, cells were switched to fresh media and were allowed to enter mitosis (∼35 minutes following drug removal). At mitotic entry, cells were collected by mitotic shake-off and were re-plated (0 hr). Time points were taken as cells exited M-phase and entered G_1_. Lysates were harvested and analyzed by immunoblotting for IRS2 and cell cycle markers.

In agreement with previous reports of mitotic phosphorylation of IRS2 by Plk1 [29], our cell cycle analysis experiments revealed that IRS2 displays a marked electrophoretic mobility shift consistent with mitotic phosphorylation. This may owe, at least in part, to Cdk1 activity given that HeLa cells released from a double thymidine block into Cdk1 inhibitor RO3306 did not display an observable shift in IRS2 mobility as compared to those released into control (DMSO) treatment (**Figure S3**). IRS2 abundance still peaked normally at this time point in the presence of RO3306, suggesting that the increase in IRS2 abundance was not dependent on Cdk1 activity. Together, these results support previous findings [29] that IRS2 is subject to cell-cycle dependent phosphorylation and that its abundance peaks in M-phase and falls in early G_1_ in multiple cell lines.

### Cdh1 control of IRS2 degradation depends on an IRS2 D-box motif

Using the SLiMSearch analysis tool [16], we found that IRS2 contains four minimal D-box motifs (RxxL), one extended D-box motif (RxxLxxxxN) and no KEN-box motifs. Of the four minimal D-box motifs, none bears strong consensus similarity to previously validated D-box motifs, and one exists in a highly structured region of the protein [24]. Because of its high SLiMSearch parameter scores (**Table 1**), we focused our efforts on determining whether the extended D-box motif located in the C-terminal third of IRS2 is required for its APC/C^Cdh1^-dependent stability. IRS2’s extended D-box (amino acids 972-980 in human IRS2) is highly conserved in placental mammals despite overall divergence in much of the C-terminus (**Figure 4A**), suggesting that this sequence likely has a conserved function.

**Figure 4:**
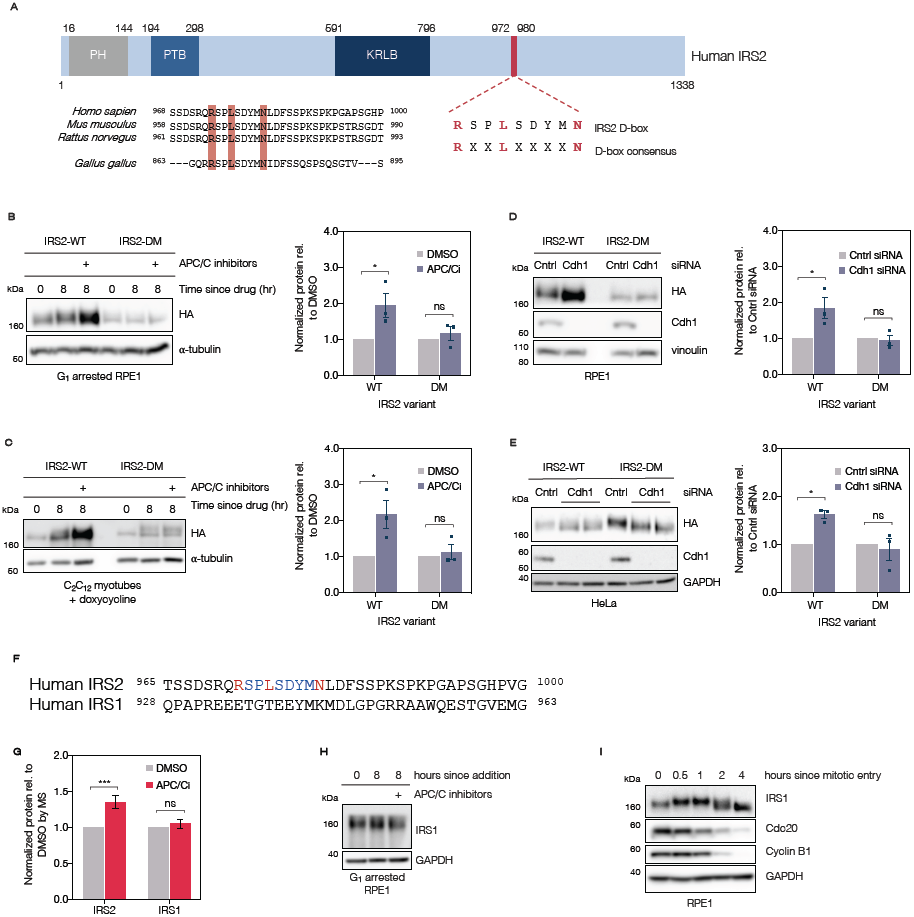
Cdh1’s ability to control IRS2 levels depends on a C-terminal D-box motif. (A) (*top*) Schematic depicting IRS2’s protein domain structure. PH = pleckstrin homology domain, PTB = phosphotyrosine binding domain, KRLB = kinase regulatory-loop binding region. IRS2’s C-terminal full D-box motif is highlighted in red. (*bottom*) Comparison of IRS2’s D-box conservation among placental mammals. (B) RPE1 cells stably expressing lentivirus-derived, doxycycline-inducible, C-terminally HA-tagged IRS2 constructs were arrested in G_1_ with 1 µM palbociclib for 20 hours in the absence of doxycycline. Following arrest, samples were either collected or DMSO or APC/C inhibitors (6 µM proTAME + 50 µM apcin) were added for an additional 8 hours. Quantification of immunoblots shown at right: HA levels were normalized to a loading control and are plotted relative to DMSO levels. Error bars = mean ± SEM. * : *p* = 0.0187; ns : *p* = 0.816 (C) C_2_C_12_ myoblasts stably expressing lentivirus-derived, doxycycline-inducible, C-terminally HA-tagged IRS2 constructs were grown to confluence and switched to low serum media supplemented with ITS (differentiation media) and doxycycline. Cells were allowed to differentiate into myotubes for three days (with media refreshment every 24 hours), at which point (0 hr) either DMSO or APC/C inhibitors (6 µM proTAME + 50 µM apcin) for an additional 8 hours in the presence of doxycycline. Quantification of immunoblots shown at right: HA levels were normalized to a loading control and are plotted relative to DMSO levels. Error bars = mean ± SEM. * : *p* = 0.0118; ns : *p* = 0.910. (D) Asynchronous RPE1 cells stably expressing lentivirus-derived, doxycycline-inducible C-terminally HA-tagged IRS2 constructs were transfected with a non-targeting (control) siRNA or an siRNA directed against Cdh1 for 24 hours in the absence of doxycycline. Quantification of immunoblots shown at right: HA levels were normalized to a loading control and are plotted relative to DMSO levels. Error bars = mean ± SEM. * : *p* = 0.0132; ns : *p* = 0.963. (E) Asynchronous HeLa cells stably expressing lentivirus-derived, N-terminally FLAG-HA tagged IRS2 constructs were transfected with a non-targeting (control) siRNA or an siRNA directed against Cdh1 for 24 hours. Quantification of immunoblots shown at right: HA levels were normalized to a loading control and are plotted relative to DMSO levels. Error bars = mean ± SEM. * : *p* = 0.0131; ns : *p* = 0.803. (F) Comparison of the Hs IRS2 D-box sequence with the corresponding region from Hs IRS1. (G) MS-quantified IRS1 and IRS2 abundance in G_1_ APC inhibitor proteomics (**Figure 1**). IRS1 abundance was quantified based on 5 peptides (4 unique) in 3 biological replicates; IRS2 was quantified based on 3 peptides (all unique) in 3 biological replicates. (H) RPE1 cells were subject to the same conditions described in **Figure 1A**, and cell lysates were analyzed by immunoblotting for IRS1 abundance (I) RPE1 cells were treated as in **Figure 3C**. Cell lysates were analyzed by immunoblotting for IRS1 abundance.

To test whether IRS2’s full D-box is relevant for its Cdh1-dependent degradation, we generated a mutant IRS2 construct bearing an R972A D-box mutation (DM), which was expected to abrogate its function as a D-box [30]. To investigate the effect of expressing the IRS2-DM construct in cells, we generated doxycycline-inducible, C-terminally HA-tagged IRS2-WT and IRS2-DM RPE1 and C_2_C_12_ cell lines. In an effort to avoid saturating the system and overexpression artifacts, we sought to express tagged IRS2 variants at low levels relative to the endogenous protein (**Figure S4A**).

Because we were able to detect the tagged proteins in the RPE1 cells without adding doxycycline, we avoided using it in this cell line. As a caveat of this approach, we observed a difference in the expression levels of the IRS2 WT and DM in RPE1 cells, possibly due to differences in lentiviral titer upon cell line generation. Still, we demonstrated that changes in transgene expression were quantifiable within the linear range for both cell lines, making comparisons between controls (DMSO or control siRNA) and APC/C inhibition (APC/C inhibitors or Cdh1 siRNA) quantifiable for each (**Figure S4B**). Using these cell lines, we found that APC/C inhibition following G_1_ arrest caused accumulation of IRS2-WT but not IRS2-DM (**Figure 4B**). The degree of accumulation of the WT protein depended on the dose of APC/C inhibitors used (**Figure S4C**).

In the C_2_C_12_ cells stably expressing C-terminally HA-tagged IRS2 variants, transgene expression was undetectable in differentiated myotubes in the absence of doxycycline (data not shown), so we selected a low dose that maintained close-to-endogenous expression levels in myoblasts (**Figure S4A**). Cells were differentiated for three days in low-serum media. Following differentiation, myotubes were switched to new media containing fresh doxycycline and either DMSO or APC/C inhibitors (the “0 hr” lane in **Figure 4C**) and were collected 8 hours later. Due to the doxycycline refreshment, myotubes exhibited a slight increase in transgene expression between 0 hours (the time of drug addition) and 8 hours (the time of collection). We observed a strong increase in levels of HA-tagged IRS2-WT in the presence of APC/C inhibitors compared to DMSO-treated myotubes. In contrast, levels of HA-tagged IRS2-DM remained constant both in the presence and absence of APC/C inhibitors (**Figure 4C**).

To further validate the Cdh1-dependence of IRS2’s D-box motif, we asked whether Cdh1 knockdown by siRNA could stabilize the IRS2-DM protein. Using asynchronous RPE1 cells stably expressing C-terminally HA-tagged IRS2-WT and IRS2-DM, we found that Cdh1 knockdown by siRNA caused an accumulation of IRS2-WT relative to control-transfected cells but not IRS2-DM (**Figure 4D**). This result was repeated in HeLa cells stably expressing N-terminally FLAG-HA-tagged IRS2-WT and IRS2-DM constructs subject to the same conditions (**Figure 4E**).

IRS1 (the other primary adaptor protein for IGF1R and IR) shares 75% sequence homology with IRS2’s N-terminus and 35% homology with its C-terminus [31] but does not share the D-box motif found in IRS2’s C-terminus (**Figure 4F**). In keeping with our hypothesis that Cdh1-mediated control of IRS2 is D-box dependent, IRS1 levels did not increase in G_1_-arrested RPE1 cells treated with APC/C inhibitors as measured by either mass spectrometry (**Figure 4G**) or immunoblot (**Figure 4H**). Furthermore, while it did display a change in electrophoretic mobility compatible with mitotic phosphorylation, unlike IRS2, it did not decrease in abundance at mitotic exit in RPE1 cells (**Figure 4I**). Taken together, the findings described above indicate that APC/C^Cdh1^ controls IRS2 levels in manner that is dependent upon its C-terminal D-box motif.

### IRS2 is required for normal expression of many proteins involved in mitosis

Many reported APC/C^Cdh1^ substrates (including several of those identified in our initial proteomics screen) are required for normal cell cycle progression. Because IR/IGF1R transduction promotes a variety of transcriptional programs [2], we hypothesized that IRS2 might promote the expression of proteins involved in cell cycle control. To investigate this, we generated two IRS2 knockout RPE1 cell lines using CRISPR/Cas9 (**Figure 5A)**, henceforth referred to as ΔIRS2-A and ΔIRS2-B. Using these cells, we again employed TMT-coupled quantitative proteomics. The proteomes of wild-type, ΔIRS2-A, and ΔIRS2-B cell lines were analyzed in biological triplicate, and relative abundances were ascertained based on TMT reporter ion signal-to-noise values. Hierarchical clustering indicated that the proteomes of the two knockout cell lines analyzed were more similar to each other than either knockout cell line was to wild-type (**Figure S5A**), suggesting that deletion of IRS2 produced similar effects in both cell lines. In order to exclude aberrancies that may have accrued during the CRISPR process or as a result of clonal expansion, we focused the scope of our analysis to proteins that changed significantly (*p* < 0.05) by more than 20% in both IRS2 knockout clones relative to the WT cell line (**Figures 5B-5C**). We found 239 proteins that decreased by >20% in both IRS2 knockout lines relative to the wild type line and 300 proteins that increased by >20% (**Figures 5B-5C, S5B**).

**Figure 5:**
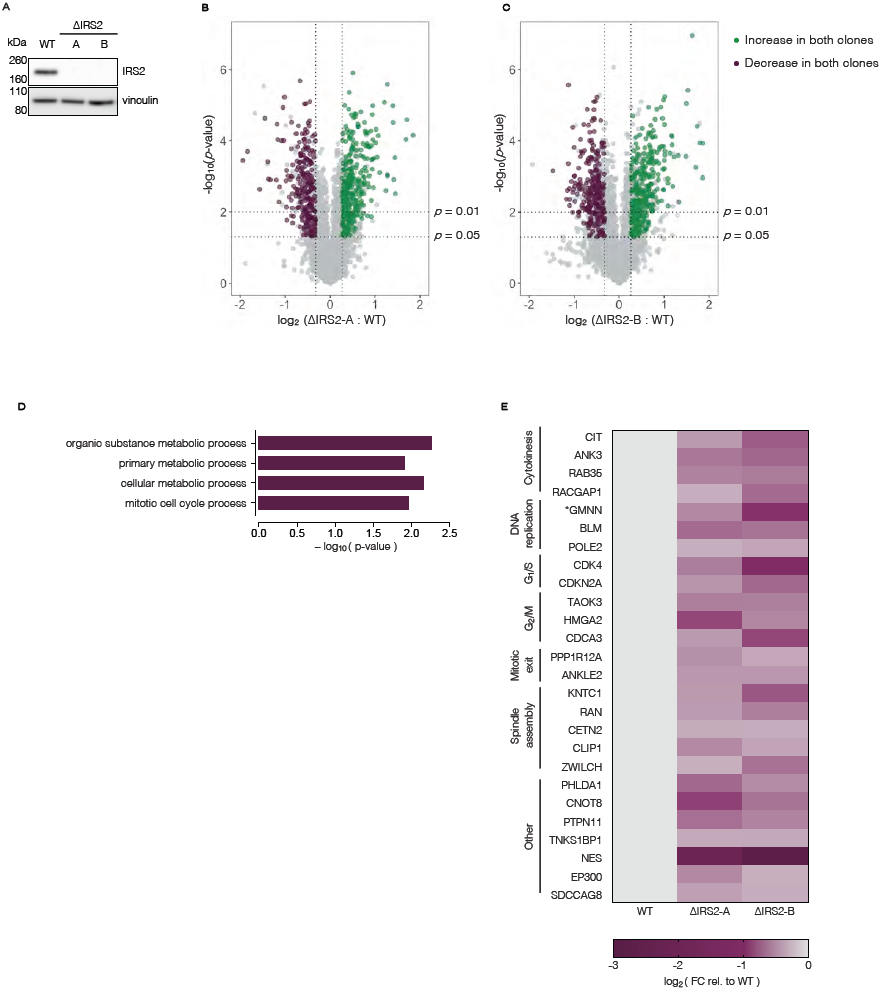
IRS2 knockout cell lines are defective in mitotic cell cycle-related protein expression. (A) WT, ΔIRS2-A, and ΔIRS2-B cell line lysates were analyzed for IRS2 expression by immunoblotting. (B-C) Volcano plots comparing proteomes of ΔIRS2 cell lines with WT cell line. Proteins that significantly decrease > 20% (*p*-value < 0.05) in both cell lines compared to wild-type are shown in purple; proteins that significantly increase > 20% (*p*-value < 0.05) in both cell lines compared to WT are shown in green. (D) Gene ontology (GO) term enrichment of proteins that decrease in both ΔIRS2 cell lines relative to WT cells. (E) Heat map depicting cell cycle-related protein abundance changes between ΔIRS2 cell lines and WT cells.

We conducted gene enrichment analysis of the proteins that increased (**Figure S5C-S5D**) or decreased (**Figure 5D**) by >20% in both knockout cell lines relative to wild type cells. Of the 239 proteins that were depleted by >20% in both knockout cell lines, we found a statistical over-representation of proteins participating in metabolic processes characteristic of IR signal transduction. Notably, we also found an over-representation of proteins involved in mitotic cell cycle regulation in this subset (**Figure 5D**). This suite of proteins included regulators of mitotic entry and exit as well as several factors involved in spindle assembly (**Figure 5E**). Importantly, IRS2 KO cells divide as the same rate as WT cells (**Figure S6A-S6B**), indicating that this downregulation is not due to a bulk loss of viability or cell cycle arrest. Consistent with the fact that strong depletion of most critical cell cycle regulators renders cells inviable, most of the observed changes in cell cycle-related genes were relatively modest (**Figure S6C**). Based on these data, we conclude that IRS2 is important for promoting the expression of a suite of proteins involved in orchestrating the mitotic cell cycle, and deletion of IRS2 stunts their expression in RPE1 cells.

### IRS2 expression promotes a functional spindle assembly checkpoint

Because many of the factors that were depleted in IRS2 knockout cell lines are involved in regulating the events of mitosis, we sought to investigate whether IRS2 KO cell lines display phenotypic differences from wild-type cells under conditions of mitotic stress—in this case, activation of the spindle assembly checkpoint (SAC). Using a high content nuclear imaging assay to measure mitotic fraction [17], we asked whether IRS2 knockout cell lines display mitotic arrest differences compared to wild-type cells when treated with spindle poisons. Wild-type cells treated with nocodazole (a microtubule destabilizing agent) arrested in mitosis in a dose-dependent manner, whereas both IRS2 knockout cell lines displayed depressed mitotic arrest (**Figure 6A** and **Figure S7A**). This was also true to a lesser extent in the presence of S-trityl-L-cysteine (STLC), an Eg5 inhibitor (**Figure 6A**). Importantly, this was not due to a reduction in the rate of mitotic entry (**Figure S6B**).

**Figure 6:**
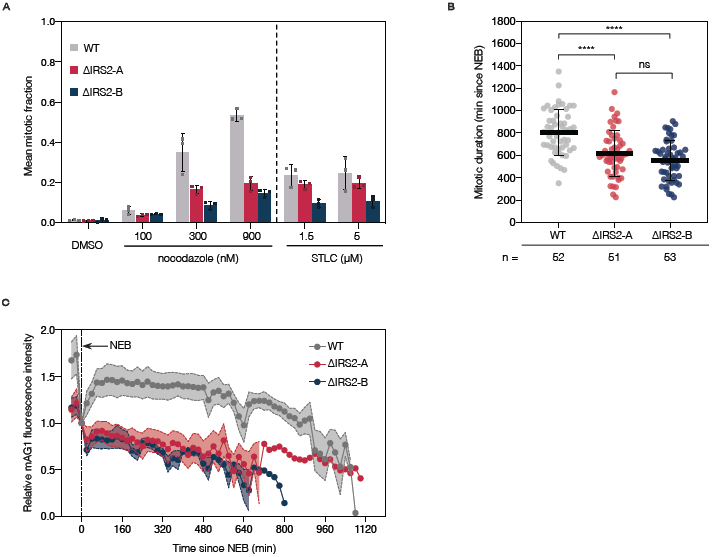
IRS2 expression promotes a functional spindle assembly checkpoint. (A) Analysis of fraction of cells in mitosis for RPE1 wild type (WT) and IRS2 KO cell lines treated with the indicated doses of nocodazole and S-trityl L-cysteine (STLC) for 18 hours. Mitotic fraction measurements were made using a high content fixed cell imaging assay based on DAPI intensity of stained nuclei. Error bars = mean ± SD. (B) Asynchronous RPE1 wild type (WT) or IRS2 KO cell lines were treated with 300 nM nocodazole and imaged every five minutes by widefield time lapse microscopy for 36 hours. Each point represents an individual cell’s mitotic duration, measured as the time from nuclear envelope breakdown (NEB) to division, slippage, or cell death. Error bars = mean± SD. *p*-values were calculated by one-way ANOVA. **** = *p* < 0.0001. ns = not statistically significant. (C) Asynchronous RPE1 wild type (WT) or IRS2 KO cell lines expressing mAG1-geminin(1-110) were treated as in (C). mAG1 fluorescence intensity was measured from nuclear envelope breakdown (NEB) until division, slippage, or cell death (n = 10 for all three cell lines). Error bars = mean ± SEM. Fluorescence intensity was background subtracted and normalized to intensity at NEB.

We further evaluated this phenotype by live-cell imaging. Consistent with the results from the fixed-cell assay, IRS2 knockout cell lines also had a significantly shorter mitotic duration compared to wild-type cells (*p* < 0.0001 in both cases) when treated with 300 nM nocodazole (**Figure 6B** and **Figure S7B**). We next analyzed the effect of IRS2 knockout on APC/C activity in cells expressing mAG1-geminin (1-101), an APC/C^Cdc20^ substrate that is stabilized by the spindle assembly checkpoint [32]. We found that wild-type cells displayed an accumulation of mAG1 fluorescence early in mitotic arrest prior to a gradual reduction due to leaky APC/C activity [33]. In contrast, both IRS2 knockout cell lines display depressed mAG1 accumulation, followed by a more rapid loss of fluorescence signal, consistent with higher APC/C^Cdc20^ activity due to a weakened checkpoint (**Figure 6C**). This phenotype, along with the shorter mitotic duration and lower mitotic fraction in the presence of spindle poisons, is consistent with cells bearing a defective mitotic spindle assembly checkpoint. Based on these data, we conclude that IRS2 expression promotes a functional spindle assembly checkpoint in RPE1 cells.

## Discussion

Based on the results of an unbiased proteomic screen, we provide evidence that IRS2, a critical mediator of IR/IGF1R signaling, is a direct APC/C^Cdh1^ substrate. We demonstrate that IRS2 is stabilized by APC/C inhibition and Cdh1 knockdown in multiple cell types and that this depends on IRS2’s C-terminal D-box motif. In contrast, we find that IRS1, a closely related IRS2 paralog that lacks a D-box, is not subject to regulation by the APC/C. Taken together, these results show that APC/C activity directly controls IRS2 levels in a D-box dependent manner.

We identified a high-mobility form of IRS2 that accumulates under APC/C inhibition, likely corresponding to a difference in phosphorylation given that IRS2 has ∼150 annotated threonine, serine, and tyrosine phosphorylation sites [34]. This suggests that IRS2’s APC/C-dependent stability could be regulated by phosphorylation, possibly at sites near or within the D-box, which is an intriguing topic for future study. Consistent with this, IRS2 phosphorylation is known to impact its stability in other contexts, including following prolonged exposure to insulin or following mTOR activation [2]. Furthermore, there is a strong precedent for phospho-regulation of APC/C degrons modulating substrate stability under specific conditions [35-37].

Many APC/C substrates are involved in cell cycle regulation, and previous studies have suggested a relationship between IRS2 and cell cycle progression. IRS2 can stimulate cell cycle entry via Cdk4 activation [38] and is important for sustaining proliferation in 32D myeloid cells and pancreatic β cells [39, 40]. Based on these findings and our identification of IRS2 as an APC/C substrate, we further investigated the role of IRS2 in regulating cell division. Proteomic analyses of RPE1 cells lacking IRS2 reveal lower expression of well-characterized cell cycle proteins compared to wild-type cells. Because these proteins are involved in critical processes like cytokinesis, DNA replication, cell cycle transitions, and spindle assembly, we investigated whether IRS2 knockout cell lines display cell cycle progression defects. We find that cells lacking IRS2 have an impaired ability to arrest following spindle assembly checkpoint activation in M-phase, thereby implicating IRS2 in promoting a functional spindle assembly checkpoint.

Despite the well-established importance of sustained IRS2 levels in many tissue types, little is known about what factors regulate its turnover. While several distinct ubiquitin ligases control IRS1 stability (Fbxw8, Cbl-b, Fbxo40, SOCS1/3, MG53, and others) [8-12], only SOCS1/3 have been implicated in the ubiquitin mediated proteolysis of IRS2 [11] until now. Thus, our work establishes APC/C^Cdh1^ as the first known ubiquitin ligase that targets IRS2 but not IRS1. Furthermore, our results suggest that APC/C^Cdh1^-mediated IRS2 degradation is relevant in broad biological contexts since we were able to demonstrate this mechanism of regulation in multiple cell lines.

Over the past several years, a number of connections between growth factor signaling and APC/C-mediated regulation have emerged. SKIL/SnoN, an APC/C substrate involved in TGFβ signaling, implicates APC/C activity in modulating the expression of TGFβ target genes [41]. Another APC/C substrate, CUEDC2, controls the stability of the progesterone receptor [42]. Regarding IR/IGF1R signaling, connections to APC/C-mediated regulation have been more opaque. Multiple reports have shown that Cdh1 interacts with PTEN, a phosphatase that antagonizes signal transduction through the IR pathway by dephosphorylating phosphoinositide-3,4,5-triphosphate (PIP_3_) [43, 44]. Others have demonstrated that components of the mitotic checkpoint complex (which inhibit APC/C^Cdc20^) potentiates IR signaling via IR endocytosis [45, 46]. Despite these links, there have been no reports of direct APC/C substrates that are involved in IR signaling until now.

Based on the data presented here, we propose a model (**Figure 7**) in which IRS2’s APC/C-mediated degradation in G_1_ serves to limit IRS2-dependent signaling during G_1_. Upon APC/C inactivation, IRS2 is able to accumulate and stimulate signaling required for normal progression through the latter stages of the cell cycle, including the expression of proteins required for mitotic spindle checkpoint function. This model is consistent with previous studies that implicate IRS2 in promoting the expression of cell cycle-related genes, including mitotic cyclins (A and B) in mouse granulosa cells [47]. Furthermore, IR signal transduction promotes the expression of Plk1 (a mitotic kinase) and CENP-A (a centromere protein) in β cells through a mechanism that appears to depend on IRS2 rather than IRS1 [40, 48].

**Figure 7:**
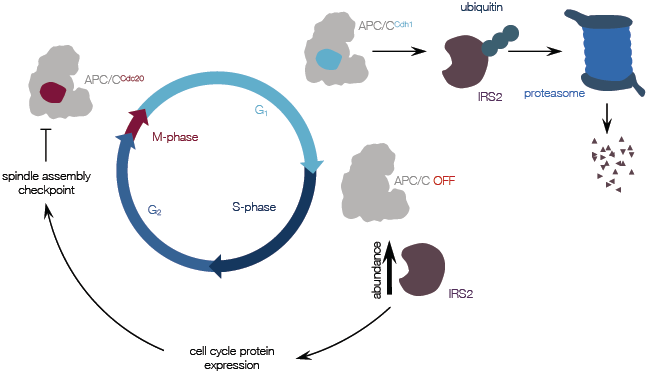
Model for IRS2’s role in cell cycle control. IRS2 is targeted for proteasomal degradation by APC/CCdh1 during G_1_. When APC/C is inactivate, IRS2 protein accumulates, potentially allowing it to stimulate the expression of cell cycle-related proteins either through IR-mediated action [48] or through another receptor tyrosine kinase. Some of the proteins that are expressed through this pathway may be required for a robust spindle assembly checkpoint, which directly inhibits APC/C^Cdc20^ during M-phase.

Our findings suggest that APC/C^Cdh1^ modulates IRS2-dependent signaling but not IRS1-dependent pathways. In IRS2-deficient mice with consequent type 2 diabetes, some have attributed the reduced β cell mass to a failure of β cells to re-enter the cell cycle following division [40]. Our findings that APC/C^Cdh1^ inhibition stabilizes IRS2 and that IRS2 promotes the expression of cell cycle regulatory proteins, coupled with data from others showing that IRS2 can stimulate cell cycle entry [38], suggest that APC/C^Cdh1^ inhibition may represent a possible approach for stimulating proliferation in quiescent β cells via the stabilization of IRS2.

## Supporting information

APC/C inhibition in G1 proteomics

Reported APC/C substrates

204 protein subset

IRS2 KO cell line proteomics

## Acknowledgements

We thank the ICCB-Longwood Screening Facility at Harvard Medical School for assistance with high content imaging assays and the Nikon Imaging Facility at Harvard Medical School for assistance with time lapse and fluorescence microscopy. We thank Kyle Copps, Pere Puigserver, and Christine Vogel for feedback on the manuscript. This work was funded by NIH grant 1R35GM127032 to R.W.K and NIH grant GM67945 to S.P.G.

## Author Contributions

S.M. designed, performed, and analyzed all experiments aimed at identifying new APC/C substrates in RPE1 cells. S.M. performed all experiments characterizing IRS2 as an APC/C substrate. S.M. generated all recombinant cell lines used in this study and the IRS2 CRISPR knockout cell lines. S.M. characterized mitotic arrest defects in IRS2 CRISPR knockout cell lines using time lapse microscopy and high-content imaging assays. S.M. prepared all samples for mass spectrometry with assistance from Q.Y.

Q.Y. and S.P.G. performed mass spectrometry analysis of G_1_ RPE1 cells treated with APC/C inhibitors and IRS2 CRISPR knockout cell lines. Q.Y. and S.P.G. provided reagents.

R.W.K. assisted with experimental design and data analysis.

S.M. and R.W.K. conceived of the project and wrote the manuscript with input from both other authors.

## Experimental Procedures

### Cell Culture and Synchronization

All cell lines used in this work (HeLa, C_2_C_12_, hTERT-RPE1-FUCCI, HEK293T) were cultured in a humidified incubator at 37ºC in the presence of 5% CO_2_. HeLa, hTERT-RPE1 and C_2_C_12_ cells were obtained from American Type Culture Collection (ATCC), and hTERT-RPE1 cells were modified with FUCCI constructs [20] with the permission of the RIKEN Institute. HeLa cells were grown in DMEM with 10% FBS. Proliferating C_2_C_12_ myoblasts were grown in DMEM with 15% FBS, whereas differentiated myotubes were cultured in differentiation media, consisting of DMEM with 2% horse serum and 1x insulin, transferrin, selenium (ITS) Premix Universal Culture Supplement (Corning, 354350). hTERT-RPE1-FUCCI cells were grown in DMEM/F12 with 10% FBS supplemented with 0.01 mg/ml hygromycin B (Corning, 30-240-CR). HEK293T cells used for lentivirus generation were a gift from Wade Harper and were cultured in DMEM with 10% FBS. All cell lines tested were negative for mycoplasma contamination (Lonza LT07-218).

HeLa cells were synchronized by double thymidine block by treating with 2 mM thymidine for 18 hours, releasing for 8 hours, and re-treating with 2 mM thymidine for 19 hours. HeLa cells synchronized by thymidine-nocodazole block were treated with 2 mM thymidine for 20 hours, released for 8 hours, then treated with 300-330 nM nocodazole for 15 hours. Mitotic cells were collected by shake-off and re-plated in drug-free media for cell cycle time course experiments.

RPE1 cells were synchronized by RO3306 treatment by treating with 7.5 µM RO3306 for 18 hours before releasing into fresh media for 30-40 minutes, after which cells were collected by mitotic shake-off and re-plated for cell cycle time course experiments. For G_1_ arrest experiments, RPE1 cells were treated with 1 µM palbociclib for 20 hours.

To differentiate C_2_C_12_ myoblasts into myotubes, cells were grown to confluence and washed 2x in DMEM with 2% horse serum before switching to differentiation media. Cells were incubated for 72 hours, with media changes every 24-36 hours. Differentiation into myotubes was monitored visually as well as by immunoblotting for MyoD, a myogenic marker.

### Immunoblotting

Cell extracts were prepared in lysis buffer (10 mM Tris HCl pH 7.4, 100 mM NaCl, 1 mM EDTA, 1 mM EGTA, 1 mM NaF, 1 mM PMSF, 20 mM Na_4_P_2_O_7_, 2 mM NA_3_VO_4_, 1% Triton X-100, 10% glycerol, 0.1% SDS, and 0.5% deoxycholate) supplemented with Pierce protease inhibitor tablets (Thermo Fisher Scientific, A32963) and Pierce phosphatase inhibitor tablets (Thermo Fisher Scientific, A32957). Pellets were incubated in lysis buffer on ice for 30 minutes with vortexing and were centrifuged at 13,000rpm for 10 minutes to clear the lysate. Protein concentrations were determined using a bicinchoninic acid (BCA) assay (Thermo Fisher Scientific, 23225). Supernatants were re-suspended in NuPAGE LDS sample buffer (Thermo Fisher Scientific, NP0008) supplemented with 100 mM dithiothreitol (DTT) and boiled at 100ºC for 5 minutes. Equal masses of lysates were separated by SDS-PAGE using either 4-12% Bis Tris gels or 3-8% Tris acetate gels (Thermo Fisher Scientific). All IRS2 immunoblots were separated on 3-8% Tris acetate gels with the exception of those shown in **Figures 5A** and **S4B**, which were separated on 4-12% Bis Tris gels. Proteins were transferred to polyvinylidene difluoride (PVDF) membranes (Thermo Fisher Scientific, 88518).

Membranes were blocked in 5% non-fat dry milk in Tris-buffered saline with 0.1% Tween (TBS-T) before incubating with primary antibodies overnight at 4ºC with agitation. Membranes were probed with secondary antibodies dissolved in 5% milk in TBS-T for 1-2 hours at room temperature before developing with an Amersham 600RGB imaging system. Quantification of immunoblots was done using ImageJ [59].

### Antibodies

The following commercially available primary antibodies were used for immunoblotting: anti-IRS2 (Cell Signaling Technologies, 4502) 1:750; anti-Cdh1/Fzr1 (Sigma Aldrich, CC43) 1:500; anti-anillin [60] 1:1000; anti-Aurora B (Bethyl, A300-431) 1:1000; anti-Eg5 (Cell Signaling Technologies, 4203) 1:1000; anti-Top2A (Cell Signaling Technologies, 12286) 1:1000; anti-TK1 (Cell Signaling Technologies, 8960) 1:1000; anti-Mps1 (Abcam, ab11108), 1:1000); anti-APC3 (BD Transduction Laboratories, 610455) 1:500; anti-cyclin B1 (Santa Cruz Biotechnology, sc-752) 1:500; anti-Cdc20 (Santa Cruz Biotechnology, sc-8358) 1:500; anti-c-Myc (9E10, Santa Cruz Biotechnology, sc-40) 1:1000; anti-HA-peroxidase (Sigma Aldrich), 1:1500; anti-cyclin A2 (Santa Cruz Biotechnology, sc-596) 1:500; anti-IRS1 (Cell Signaling Technologies, 2382) 1:750; anti-MyoD1 (Cell Signaling Technologies, 13812) 1:750; anti-GAPDH (Abcam, ab8245) 1:2000; anti-α tubulin (Abcam, ab7291 and Santa Cruz Biotechnology, sc-8035) 1:1000 for both; anti-vinculin (Santa Cruz Biotechnology, sc-73614) 1:2000. Secondary antibodies used: anti-rabbit IgG-HRP (GE Healthcare, NA934) and anti-mouse IgG-HRP (GE Healthcare, NA931V), both at 1:3000 dilutions.

### Compounds

The following chemicals were used: palbociclib (LC Laboratories, P-7722), proTAME (Boston Biochem, I-440), MG132 (474790, Calbiochem), S-trityl L-cysteine (STLC, Alfa Aesar, L14384), thymidine (Sigma Aldrich, T9250), nocodazole (Sigma Aldrich, 31430-18-9), RO3306 (AdipoGen Life Sciences, AGCR13515M), doxycycline hyclate (Sigma Aldrich, D9891). Apcin was custom synthesized by Sundia MediTech Company (Lot #A0218-10069-031) using methods described previously [17]. All compounds were dissolved in dimethyl sulfoxide (DMSO), with the exception of thymidine and doxycycline, which were dissolved in Dulbecco’s phosphate buffered saline (DPBS, Corning, 21-030-CV). Dissolved compounds were stored at −20°C prior to use.

### CRISPR/Cas9 mediated gene editing

A TrueGuide crRNA directed against exon 1 of Hs IRS2’s coding region (target DNA sequence: 5’-TCG AGA GCG ATC ACC CGT TT −3’, Assay ID number: CRISPR850215_CR, Thermo Fisher Scientific) was annealed to the TrueGuide tracrRNA (Thermo Fisher Scientific, A35507) according to manufacturer protocol. hTERT RPE1-FUCCI cells were co-transfected with TrueCut Cas9 protein v2 (Thermo Fisher Scientific, A36496) and the annealed tracrRNA:crRNA complex using the Lipofectamine CRISPRMAX Cas9 Transfection reagent (Thermo Fisher Scientific, CMAX00003) according to manufacturer protocol. Transfected cells were incubated for two days before switching to fresh media and expanding. Single cell clones were isolated using the limiting dilution method in a 96-well format, and clonal cell lines were expanded before screening for knockouts by immunoblotting.

### Site directed mutagenesis

R777-E111 Hs.IRS2 and R777-E111 Hs.IRS2-nostop were gifts from Dominic Espositio (Addgene plasmid #70395 and #70396, respectively). Both of these plasmids encode codon optimized sequences for IRS2, with and without a stop codon respectively. R972A mutations were introduced into the aforementioned IRS2 clones using the Q5 Site-Directed Mutagenesis Kit (New England BioLabs) with the primers 5’ - AGA TTA TAT GAA TAA GTC CAC TGT CAG ATT ATA TG - 3’ and 5’ - GAC AGT GGA CTT GCC TGG CGA GAG TCT GAA CT - 3’ according to the manufacturer’s protocol. For N-terminally FLAG-HA-tagged constructs, the insert from R77-E111 Hs.IRS2 (WT or) was cloned into the pHAGE-FLAG-HA-NTAP vector (a gift from Wade Harper) using the Gateway LR Clonase II system (Invitrogen). For doxycycline-inducible, C-terminally HA-tagged constructs, the insert from R77-E111 Hs.IRS2-nostop (WT or DM) was cloned into pINDUCER20 (a gift from Stephen Elledge, Addgene plasmid #44012) using the Gateway LR Clonase II system (Invitrogen). The DM mutation was verified both before and after Gateway cloning by Sanger sequencing.

### Lentivirus construction

To construct lentiviruses, HEK293T cells were co-transfected with pPAX2, pMD2, and either pINDUCER-20-IRS2 or pHAGE-FLAG-HA-NTAP-IRS2 in a 4:2:1 DNA ratio using Lipofectamine 3000 (Invitrogen, L3000001) according to manufacturer’s instructions. pPAX2 and pMD2 were gifts from Wade Harper. 24 hours after transfection, HEK293T cells were switched to fresh media (DMEM + 10% FBS). 48 hours after transfection lentiviruses were harvested by clearing debris by centrifugation at 960*xg* for 5 minutes and filtering through 0.45 µm SFCA filters. Lentiviruses were either used immediately or flash frozen in liquid nitrogen and stored at −80ºC for later use.

### Stable cell line construction

To generate stable cell lines, plated HeLa, RPE1, or C_2_C_12_ cells were incubated with lentiviruses and 2 µg/ml protamine sulfate. 24 hours after viral infection, cells were switched to fresh media. 48 hours after viral infection, antibiotics were introduced. For lentiviruses derived from pINDUCER20, geneticin (Invitrogen, 10131027) was used at a concentration of 750 µg/ml for both RPE1 and C_2_C_12_ for 6-7 days. For lentiviruses derived from pHAGE-FLAG-HA-NTAP, puromycin (Sigma Aldrich, P8833) was used at a concentration of 0.5 µg/ml for 3 days. Antibiotic-selected populations of cells were expanded and used for further experiments without clonal selection.

### Small interfering RNAs (siRNAs)

Cells were transfected using RNAiMax (Invitrogen, 13778100) according to manufacturer’s instructions with the following siRNAs: siGENOME Non-Targeting Control siRNA #5 (D-001210-05, Dharmacon); ON-TARGETplus Human FZR1 siRNA (J-015377-08, Dharmacon), 25 nM; SMARTpool ON-TARGETplus Mouse Fzr1 siRNA (L-065289-01-0005), 25 nM. Cells were treated with siRNAs for 24 hours for all experiments. For experiments involving subsequent compound treatment, cells were switched to fresh media prior to the addition of compounds.

### Plasmid transfection

C_2_C_12_ myoblasts were transfected with a plasmid containing the N-terminal 88 amino acids of human cyclin B1 fused to EGFP using Lipofectamine 3000 (Invitrogen, L3000001) with the P3000 reagent according to manufacturer’s instructions. Growth media was refreshed to remove transfection reagents 24 hours post-transfection, and cells were switched to differentiation media for an additional 3 days.

For ubiquitination studies, HeLa cells were transfected with equal amounts of pCI-6xHis-hUb and/or pHAGE-NTAP-IRS2 using Lipofectamine 3000 with the P3000 reagent according to manufacturer’s instructions. For myc-Cdh1 overexpression studies, HeLa cells were transfected with 0, 0.5, 1, and 2 μg of pCS2+ myc-hCdh1 using Lipofectamine 3000 with the P3000 reagent according to manufacturer’s instructions.

### In vivo ubiquitination assay

HeLa cells expressing pCI-6xHis-hUb and pHAGE-NTAP-IRS2 were treated with 10 μM MG132 either alone or in the presence of APC/C inhibitors (6 μM proTAME + 50 μM apcin) for 8 hours. Following drug treatment, cells were collected, washed in Dulbecco’s Phosphate Buffered Saline with calcium and magnesium (Corning, 21-030-CM), and flash frozen in liquid nitrogen. Cell pellets were lysed in denaturing lysis buffer (8M urea; 100 mM Na_2_HPO_4_; 0.05% Tween-20; 10 mM imidazole HCl, pH 8.0; 100 mM Tris HCl, pH 8.0; 1x protease inhibitor cocktail) by periodic vortexing on ice. Lysates were cleared at 13,000 rpm for 10 minutes. Protein concentrations were measured by BCA assay. Equal amounts of protein were incubated with Ni-NTA agarose resin (Qiagen, 30210) for four hours, rotating at 4°C. Resins were washed three times in denaturing wash buffer (8M urea; 100 mM Na_2_HPO_4_; 0.05% Tween-20; 20 mM imidazole HCl, pH 8.0; 100 mM Tris HCl, pH 8.0) followed by three washes in native wash buffer (100 mM Na_2_HPO_4_; 0.05% Tween-20; 20 mM imidazole HCl, pH 8.0; 100 mM Tris HCl, pH 8.0). His-ubiquitin conjugates were eluted by boiling resin in 2x LDS sample buffer (Thermo Fisher Scientific, NP0008) supplemented with 200 mM DTT and 200 mM imidazole HCl, pH 8.0 for 10 minutes. Input samples were resolved on 4-12% Bis-Tris gels (Thermo Fisher Scientific), and Ni-NTA elution samples were resolved on 3-8% Tris acetate gels (Thermo Fisher Scientific). Ponceau staining was used as a loading control.

### Time lapse and fluorescence microscopy

Cells were plated in a 24-well coverslip-bottom plate (Greiner BioOne, 662892). After 24 hours, cells were treated with the indicated compounds and were imaged immediately afterwards. Plates were inserted into a covered cage microscope incubator (OkoLab) with temperature and humidity control at 37ºC and 5% CO_2_ and mounted on a motorized microscope stage (Prior ProScan III). All images were collected on a Nikon Ti motorized inverted microscope equipped with a 20x/0.75 NA Plan Apo objective lens and the Perfect Focus system. mCherry fluorescence was excited with a Lumencor Spectra-X using a 555/25 excitation filter and a 605/52 emission filter (Chroma). mAG1 fluorescence was excited using a 490/20 excitation filter and a 525/36 emission filter (Chroma). Both configurations used a Sedat Quad dichroic (Chroma). Images were acquired with a Hamamatsu Orca-R2 or Hamamatsu Flash 4.0 V2 controlled with Nikon Elements image acquisition software. Three fields of view were collected per condition, and phase contrast and/or fluorescence images were captured at 5- to 8-minute intervals (depending upon the experiment) for 24-48 hours.

Videos were analyzed using ImageJ. Mitotic duration was defined as the time from nuclear envelope breakdown (NEB) until division, death (cytoplasmic blebbing), or mitotic slippage. mAG1 and mCherry intensities were quantified manually by measuring the maximum intensity of signal for each cell in a given frame across multiple time points. For experiments analyzing fluorescence intensity during G_1_ arrest, measurements were made for all cells in a frame for each time point.

### Experimental design and statistical rationale

For G_1_ proteomics experiments, cells were treated with 1 μM palbociclib for 20 hours at which point cells were either collection (t_0_) or treated with DMSO (control) or APC/C inhibitors (6 μM proTAME + 50 μM apcin) for an additional 8 hours prior to collection (n=3). For IRS2 knockout proteomic analysis, the proteomes of one control cell line (which had been subject to the CRISPR process but expressed wild-type levels of IRS2) and two distinct clonal IRS2 knockout lines were measured (n=3). For both G_1_ APC/C inhibitor proteomics and IRS2 knockout cell line analysis, samples were analyzed in biological triplicate (e.g. from separate starter cultures) to increase statistical power. Technical replicates were not measured for either experiment. No samples were excluded from either analysis. Peptide spectral matches were filtered with a linear discriminant analysis (LDA) method to a 1% FDR [61] and a protein-level FDR of 1% was also implemented [62]. For both datasets, comparisons were calculated using a two-tailed, unpaired Student’s t-test. For the G_1_ proteomics experiment, we considered significantly changing proteins as those with a *p*-value<0.05 that increased ≥1.15-fold (based on the fold-changes observed for previously reported APC/C substrates within our dataset). For the IRS2 knockout cell line analysis, we considered significantly changing proteins as those with a *p*-value<0.05 and a <20% change in abundance. For experiments regarding the stability of IRS2-WT and IRS2-DM, *p*-values were calculated by two-way ANOVA. For fluorescence microscopy experiments that quantify mAG1 intensity in response to drug treatment over time, *p*-values were calculated by two-way ANOVA. For microscopy experiments that quantify mitotic duration following nocodazole treatment, *p*-values were calculated by one-way ANOVA. For time lapse microscopy data, *p*-values were calculated by one-way ANOVA. Gene enrichment was calculated using the AmiGO 2 search tool [63]. Error bars represent standard deviation (SD) or standard error of the mean (SEM) where indicated.

### TMT mass spectrometry sample preparation

Cells were cultured as described in biological triplicate. Cells pellets were re-suspended in urea lysis buffer: 8M urea, 200 mM EPPS pH 8.0, Pierce protease inhibitor tablets (Thermo Fisher Scientific, A32963), and Pierce phosphatase inhibitor tablets (Thermo Fisher Scientific, A32957). Lysates were passed through a 21-gauge needle 20 times, and protein concentrations were measured by BCA assay (Thermo Fisher Scientific). 100 µg of protein were reduced with 5 mM tris-2-carboxyethyl-phosphine (TCEP) at room temperature for 15 minutes, alkylated with 10 mM iodoacetamide at room temperature for 30 minutes in the dark, and were further reduced with 15 mM DTT for 15 minutes at room temperature. Proteins were precipitated using a methanol/chloroform extraction. Pelleted proteins were resuspended in 100 µL 200 mM EPPS, pH 8.0. LysC (Wako, 125-05061) was added at a 1:50 enzyme:protein ratio, and samples were incubated overnight at room temperature with agitation. Following overnight incubation, trypsin (Promega, V5111) was added at a 1:100 enzyme:protein ratio, and samples were incubated for an additional 6 hours at 37ºC. Tryptic digestion was halted by the addition of acetonitrile (ACN). Tandem mass tag (TMT) isobaric reagents (Thermo Fisher Scientific, 90406) were dissolved in anhydrous ACN to a final concentration of 20 mg/mL, of which a unique TMT label was added at a 2:1 label:peptide ratio. Peptides were incubated at room temperature for one hour with vortexing after 30 minutes. TMT labeling reactions were quenched by the addition of 10 µL of 5% hydroxylamine. Equal amounts of each sample were combined at a 1:1 ratio across all channels and lyophilized by vacuum centrifugation. Samples were re-suspended in 1% formic acid (FA)/99% water and were desalted using a 50 mg 1cc SepPak C18 cartridge (Waters, WAT054955) under vacuum. Peptides were eluted with 70% ACN/1% FA and lyophilized to dryness by vacuum centrifugation. The combined peptides were fractionated with basic pH reversed-phase (BPRP) HPLC, collected in a 96-well format and consolidated to a final of 24 fractions, out of which only alternating fractions (a total of 12) were analyzed [64]. Each fraction was desalted via StageTip, lyophilized to dryness by vacuum centrifugation, and reconstituted in 5% ACN/5% FA for LC-MS/MS processing.

### TMT mass spectrometry analysis

Data for the G_1_ APC/C inhibition experiment were collected on an Orbitrap Fusion mass spectrometer coupled to a Proxeon EASY-nLC 1000 liquid chromatography (LC) pump (Thermo Fisher Scientific), whereas data for IRS2 knockout cell line analysis were collected on an Orbitrap Fusion Lumos mass spectrometer coupled to a Proxeon EASY-nLC 1200 liquid chromatography (LC) pump. The 100 μm capillary column was packed with 30 cm of Accucore 150 resin (2.6 μm, 150Å; Thermo Fisher Scientific). Mobile phases were 5% ACN, 0.125% FA (Buffer A) and 95% ACN, 0.125% FA (Buffer B). Peptides from G_1_ APC/C inhibition experiment were separated using a 2.5 h gradient from 4% to 26% Buffer B and analyzed with a SPS-MS3 method [65]. Peptides from IRS2 knockout cell line analysis were separated using a 2 h gradient from 4% to 30% Buffer B and analyzed with a real-time search strategy [66, 67]. The mass spectrometry proteomics data have been deposited to the ProteomeXchange Consortium via the PRIDE [68] partner repository with the dataset identifier PXD018329 and 10.6019/PXD018329.

Raw data were converted to mzXML format using a modified version of RawFileReader (https://planetorbitrap.com/rawfilereader) and searched with SEQUEST [69] using an in-house proteomic pipeline against a human protein target-decoy database containing both SwissProt and TrEMBL entries (downloaded February 2014). The database was concatenated onto a database of common contaminants (e.g. trypsin, human keratins). Searches were performed with a 50 ppm precursor mass tolerance, 0.9 Da fragment mass tolerance, trypsin digest with up to 2 missed cleavages. Allowed modifications include cysteine carboxyamidomethylation (+57.02146), static TMT on lysine and peptide N-temini (+229.16293) and up to 3 variable methionine oxidation (+15.99491). Peptide spectral matches were filtered with a linear discriminant analysis (LDA) method to a 1% FDR [61] and a protein-level FDR of 1% was also implemented [62]. For peptide quantification, we extracted the TMT signal-to-noise and column normalized each channel to correct for equal protein loading. Peptide spectral matches with summed signal-to-noise less than 100 were excluded from final result. Lastly, each protein was scaled such that the summed signal-to-noise for that protein across all channels equals 100, thereby generating a relative abundance (RA) measurement.

### High content mitotic fraction assay

Asynchronous hTERT RPE1-FUCCI wild-type or IRS2 KO cell lines were plated in a black, clear-bottom 96-well plate (Corning, 3606). Plates were sealed with breathable white rayon sealing tape (Nunc, 241205) to prevent evaporation following plating and during all subsequent incubations. In experiments involving RNAi, cells were treated with siRNAs for 24 hours. Cells were switched to fresh media, and compounds were added at the indicated concentrations for an additional 18 hours. Following compound treatment, cells were fixed and stained directly without additional washing steps (to avoid the loss of loosely attached mitotic cells) with 10% formalin, 0.33 µg/mL Hoechst 33342, and 0.1% Triton X-100 in DPBS. Plates were sealed with aluminum tape (Nunc, 276014) and were incubated for 45 minutes room temperature in the dark before imaging. All experimental conditions were represented in triplicate on the same plate. Plates were imaged using an ImageXpress Micro high-content microscope (Molecular Devices) equipped with a 10x objective lens. Four images were acquired per well, yielding a total of 12 images per conditions. Images were processed automatically in ImageJ to identify and count nuclei as well as measure their maximum fluorescence intensity. ImageJ output files were pooled, and cumulative frequency curves for the maximum intensity of the cell population in each condition were computed using MATLAB. An intensity threshold was set based on the intensity of mitotic cells in control (DMSO-treated) wells to delineate interphase cells from mitotic cells. The fraction of mitotic cells was calculated as the fraction of cells above the set intensity threshold in MATLAB[17].

### Materials Availability

All mass spectrometry raw files will be available through the PRIDE archive upon publication. All other data are available in the associated supplementary data files. Further information and requests for resources and reagents should be directed to the Lead Contact, Randy King.

## Supplemental Information Legends

***Figure S1***: *Related to Figure 1*

(A) Asynchronous RPE1-FUCCI cells were treated with 1 µM palbociclib and imaged by fluorescence time lapse microscopy for 20 hours. Frames at 0, 10, and 20 hours are shown.

(B) From the experiment shown in Figure S1B, FITC intensity was quantified at 0 hours (time of drug addition) and 20 hours. Each point represents the maximum FITC intensity of an individual cell at the given time point. ns : not significant ; **** : *p*<0.0001.

(C) Asynchronous RPE1-FUCCI cells were treated with 1 μM palbociclib for 20 hours. Following G_1_ arrest, cells were treated with either DMSO or APC/C inhibitors (6 µM proTAME + 50 µM apcin) and imaged by fluorescence widefield time lapse microscopy for an additional eight hours.

(D) Quantification of the experiment shown in S1D, as explained in S1B. Error bars = SD among all of the cells quantified for each condition. ns: not significant ; **** : *p*<0.0001.

(E) RPE1-FUCCI cells were treated with 1 μM palbociclib for 20 hours. Following G_1_ arrest, cells were either switched to fresh media containing no drug (“wash out”), fresh media containing the same concentration of palbociclib (“palbociclib”), or fresh media containing the same concentration of palbociclib and APC/C inhibitors (palbociclib + APC/Ci). mAG1 intensity was quantified for all cells in a given frame every 2.7 hours for 16 hours. Each data point represents the mean intensity ± SEM for at least 90 cells from two different frames for each condition.

***Figure S2***: *Related to Figure 2B*

(A) Asynchronous C_2_C_12_ myoblasts and 3-day differentiated C_2_C_12_ myotubes were lysed and MyoD levels were measured by immunoblotting.

(B) Phase-contrast images of asynchronous (Day 0) C_2_C_12_ myoblasts and 3-day differentiated C_2_C_12_ myotubes.

(C) Asynchronous C_2_C_12_ myoblasts were transfected with a plasmid coding for the N-terminal fragment of cyclin B1 (amino acids 1-88) fused to EGFP for 24 hours. Following transfection, cells were switched to low-serum differentiation media containing ITS for three days with media refreshment every 24 hours. After 3 days, myotubes were acutely treated with either DMSO or APC/C inhibitors (6 µM proTAME + 50 µM apcin) for an additional 8 hours. Myotubes were then harvested, and lysates were analyzed for transgene expression by immunoblot.

(D) Asynchronous C_2_C_12_ myoblasts were transfected with an siRNA directed against the mouse Cdh1 sequence for 18 hours before being maintained in fresh media for an additional 18 hours. Cells were harvested, and lysates were analyzed for the expression of the indicated proteins.

***Figure S3***: *Related to Figure 3*

HeLa cells were synchronized by double thymidine block and released into S-phase either in the presence of DMSO or 5 µM RO3306. Cells were harvested at the indicated time points for analysis of the given protein abundances and phosphorylation patterns in lysate by immunoblot.

***Figure S4***: *Related to Figure 4*

(A) (*top*) Asynchronous RPE1 cells expressing doxycycline-inducible, C-terminally HA tagged IRS2 variants were treated with a dose range of doxycycline. HA and IRS2 expression levels were analyzed by immunoblotting cell lysates. Red = doxycycline dose used for all experiments. (*bottom*) Asynchronous C_2_C_12_ cells expressing doxycycline-inducible, C-terminally HA tagged IRS2 variants were treated with a dose range of doxycycline. HA and IRS2 expression levels were analyzed by immunoblotting cell lysates. Red = doxycycline dose used for all experiments.

(B) (*left*) Asynchronous RPE1 cells expressing doxycycline-inducible, C-terminally HA-tagged variants were lysed, and a dilution series of lysate was analyzed for HA expression by immunoblot. The protein amount highlighted in red, 15 μg, was the amount loaded for all immunoblot experiments. The corresponding Coomassie stained gel is shown below. (*right*) Quantification of the experiment at left. Linear regression calculations were made, and lines were plotted over data. For WT, R^2^= 0.97. For DM, R^2^=0.99

(C) *(left*) RPE1 cells stably expressing lentivirus generated, C-terminally HA-tagged IRS2 wild type (WT) and R972A (DM) constructs were arrested in G_1_ with 1 µM palbociclib for 20 hours. Cells were then acutely treated with either DMSO or the indicated dose range of APC/C inhibitors for an additional 8 hours. Cells were then harvested, and lysate was analyzed for HA expression by immunoblot. The lane denoted t_0_ indicates a sample that was collected at the time of drug addition. (*right*) Quantification of the experiment at left. Plot shows HA intensity normalized to a loading control (either GAPDH or Ponceau) and to the DMSO condition. Error bars = mean ± SEM.

***Figure S5***: *Related to Figure 5*

(A) Hierarchical clustering for the nine conditions analyzed by TMT-coupled quantitative mass spectrometry in wild type and ΔIRS2 cell lines.

(B) Venn diagrams depicting proteins that (*left*) decrease significantly >20% relative to WT cells in both ΔIRS2 cell lines and (*right*) increase significantly >20% relative to WT in both ΔIRS2 cell lines.

(C) Gene ontology (GO) term enrichment of proteins that increase in both ΔIRS2 cell lines relative to WT cells.

(D) Heat map for proteins in the “oxidation-reduction process” GO category increasing in both ΔIRS2 cell lines relative to WT cells.

***Figure S6***: Related to Figure 5

(E) Growth curves for IRS2 knockout cell lines.

(F) Asynchronous RPE1 WT and IRS2 KO cells were imaged every five minutes by widefield time lapse microscopy for 36 hours. The fraction of cells that entered mitosis over this time span was measured manually, and cumulative frequency curves for mitotic entry were plotted. WT: n= 33; ΔIRS2-A: n=42; ΔIRS2-B: n=42.

(G) Fractional abundance of cell-cycle related proteins shown in **Figure 5D-5E** depleted in ΔIRS2 cell lines relative to WT cells. ΔIRS2-A median abundance = 0.70; ΔIRS2-B median abundance = 0.66.

***Figure S7***: *Related to Figure 6*

(A) Representative frames from high-content nuclear imaging experiment for mitotic fraction based on DAPI intensity. Asynchronous RPE1 WT or IRS2 KO cell lines were treated with 900 nM nocodazole for 18 hours before fixing and DAPI staining.

(B) Asynchronous RPE1 WT and IRS2 KO cell lines were imaged every five minutes by widefield time lapse microscopy for 36 hours. Each point represents an individual cell’s mitotic duration, measured as the time from nuclear envelope breakdown (NEB) to division, slippage, or cell death. Error bars = mean ± SD. *p*-values were calculated using one-way ANOVA. ns = not statistically significant.

(C) Asynchronous RPE1 WT and IRS2 KO cell lines were imaged as in (B). Images are representative of an unperturbed mitosis from NEB to cytokinesis for each cell line. Time (in minutes) in upper right corner indicates time since NEB.

***Figure S8***: *Related to Figures 1-2*

Extended immunoblots from Figures 1 and 2

***Figure S9***: *Related to Figures 3*

Extended immunoblots from Figure 3

***Figure S10***: *Related to Figures 4-5*

Extended immunoblots from Figures 4 and 5

***Table S1***: APC/C inhibition in G_1_ proteomics

***Table S2***: Reported APC/C substrates identified by proteomics

***Table S3***: 204 protein subset

***Table S4***: IRS2 knockout cell proteomics

