## Supplemental Figures 1-10 for "The insulin receptor adaptor IRS2 is an APC/C substrate that promotes cell cycle protein expression and a robust spindle assembly checkpoint"

Supplemental Figure 1

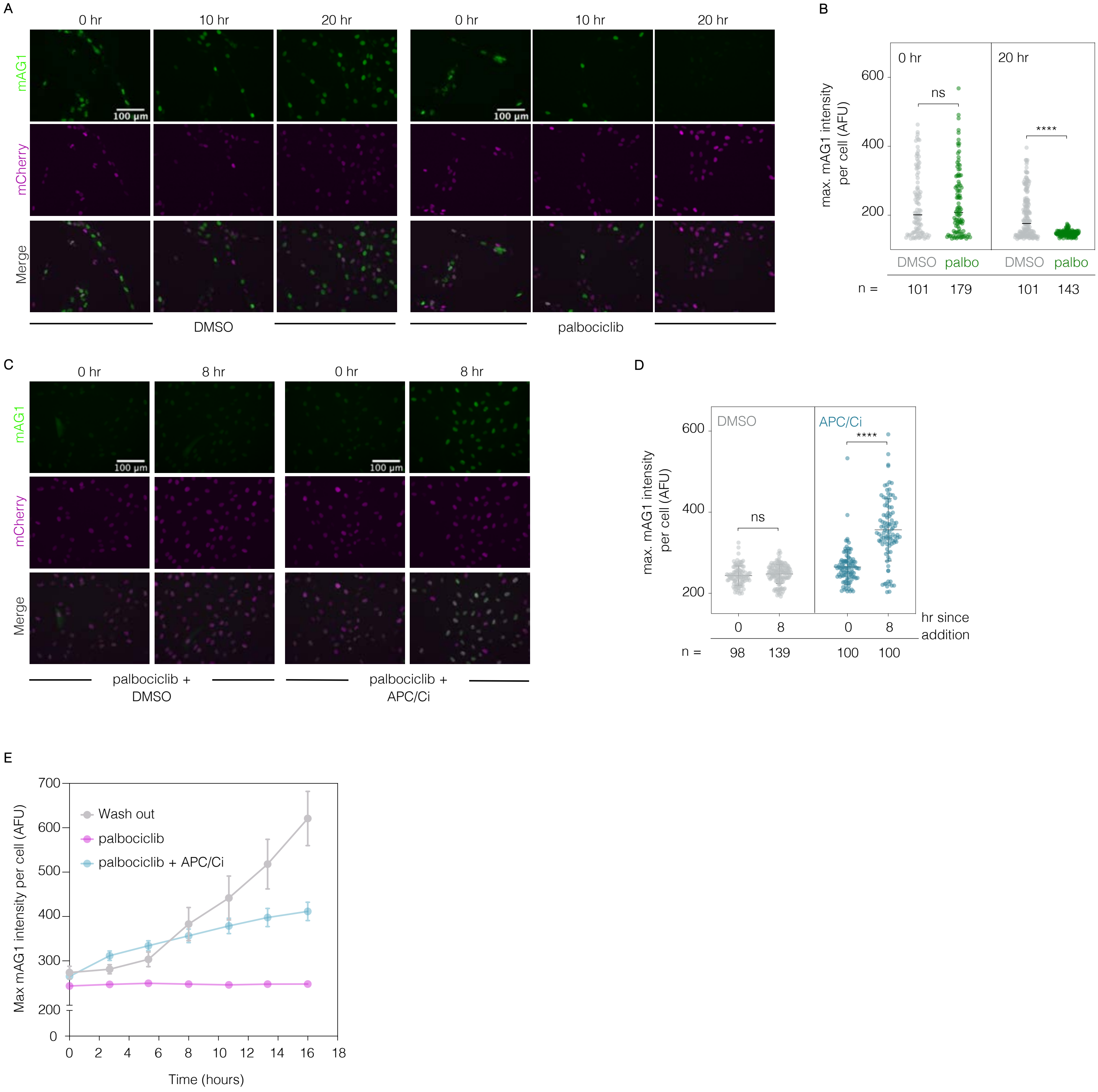

Supplemental Figure 2

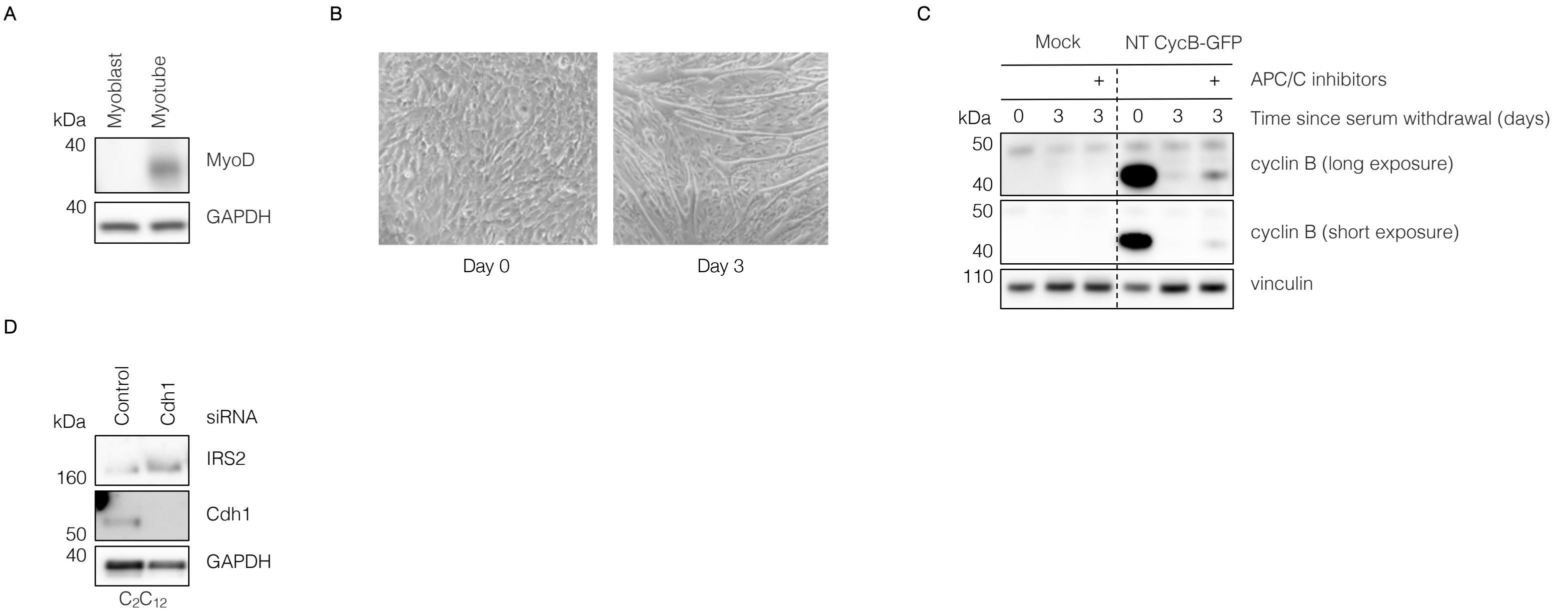

Supplemental Figure 3

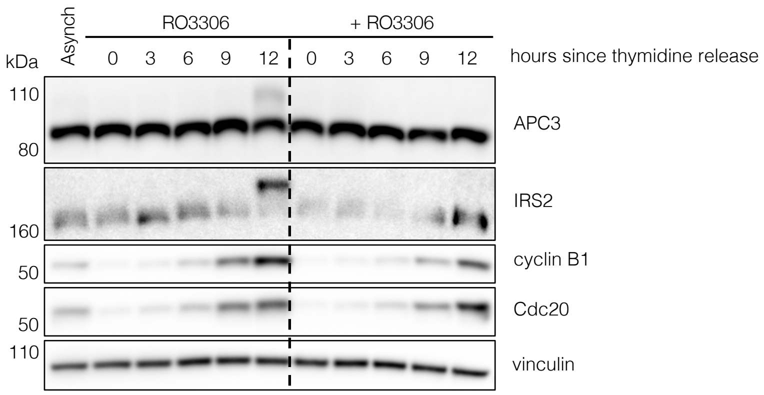

Supplemental Figure 4

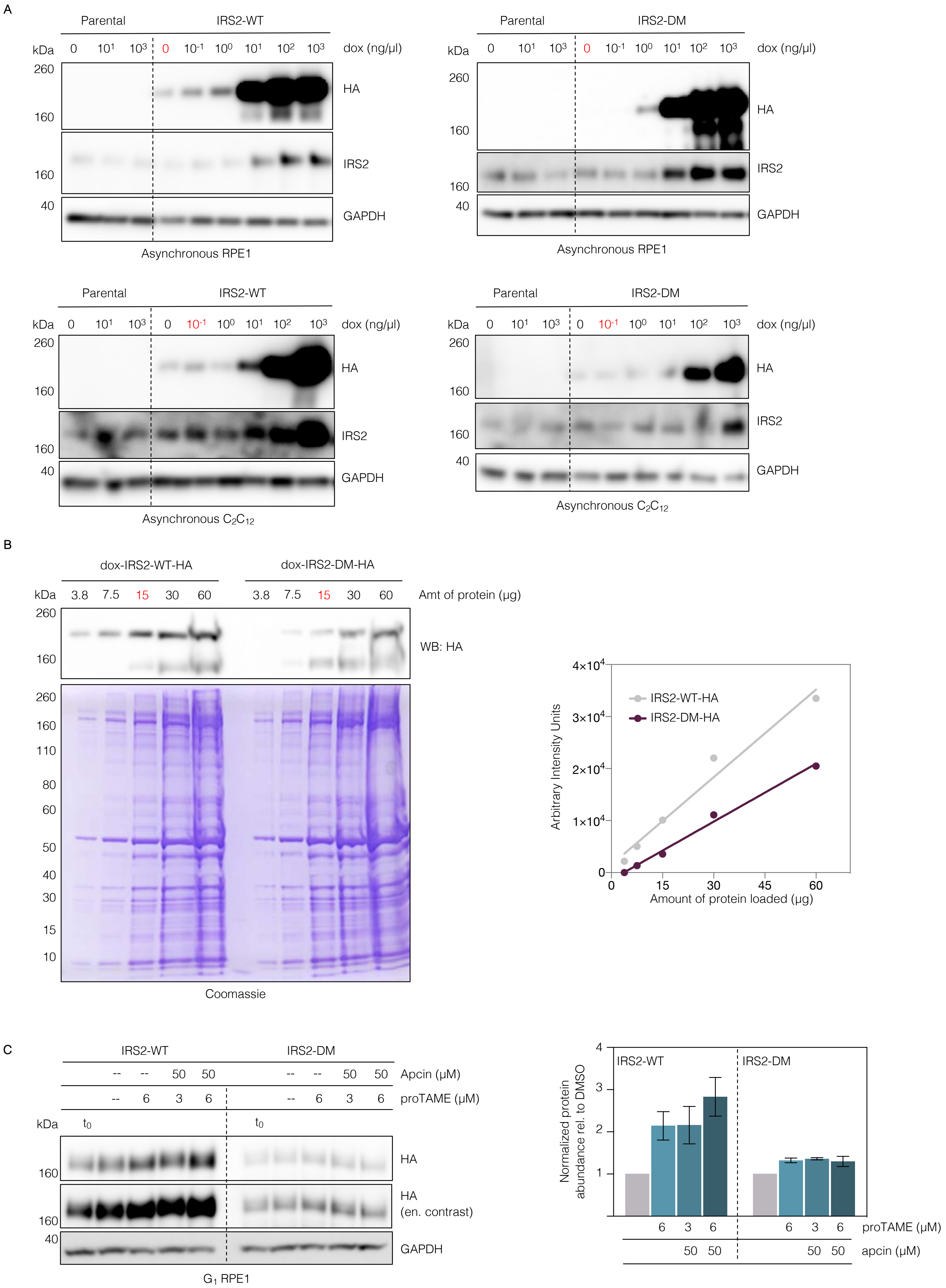

Supplemental Figure 5

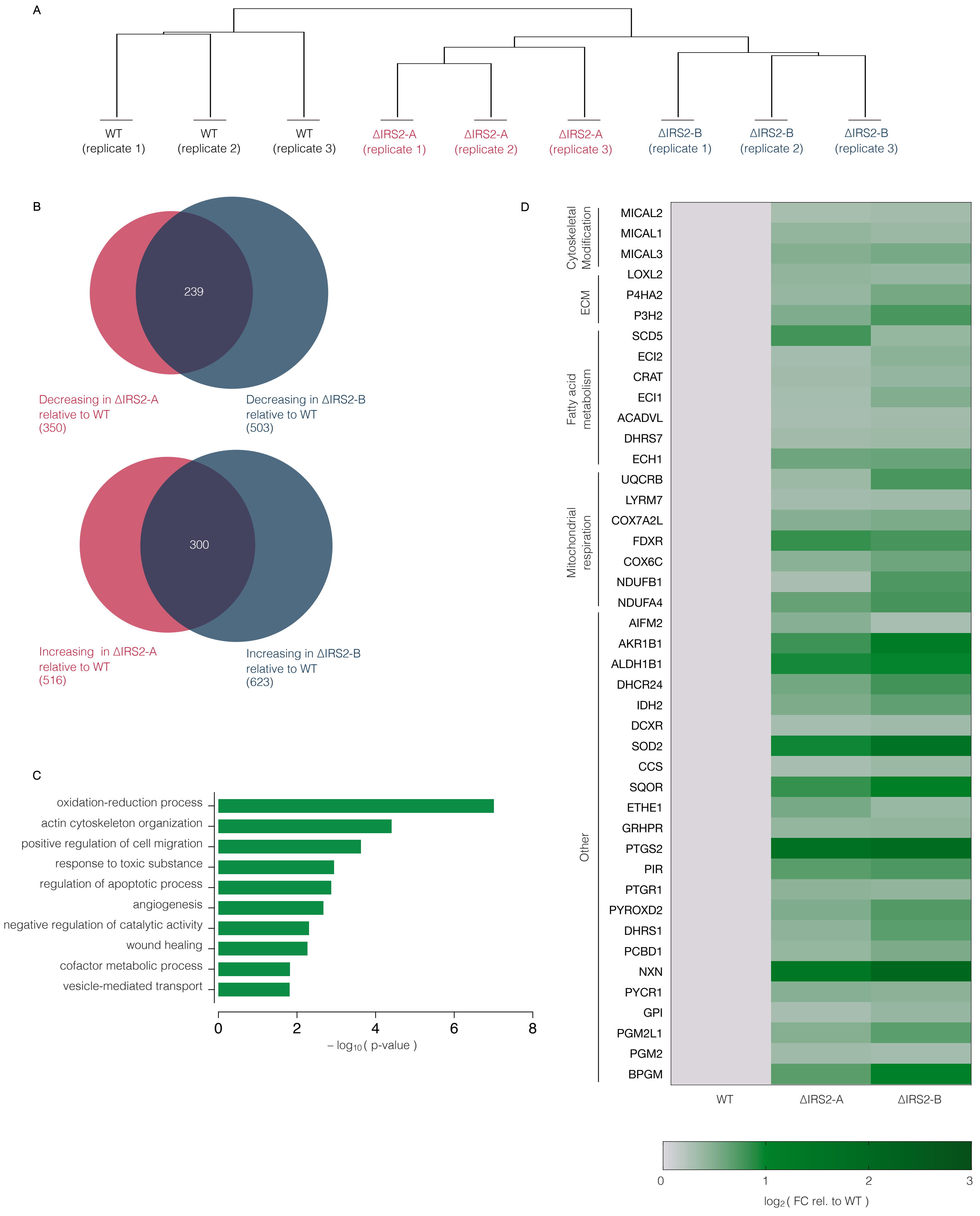

Supplemental Figure 6

A

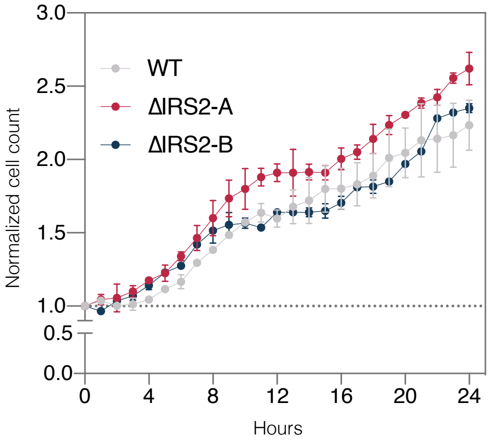


B

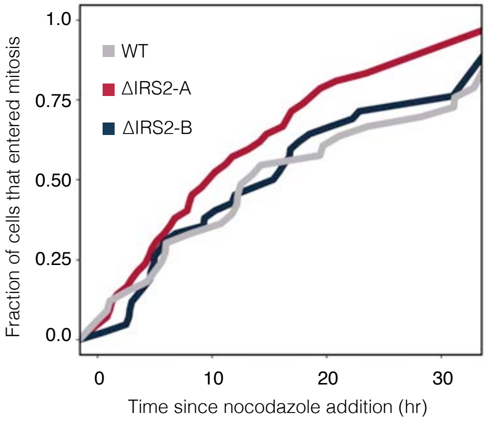


C

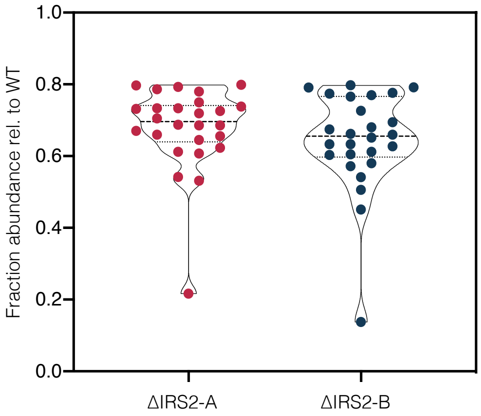

Supplemental Figure 7

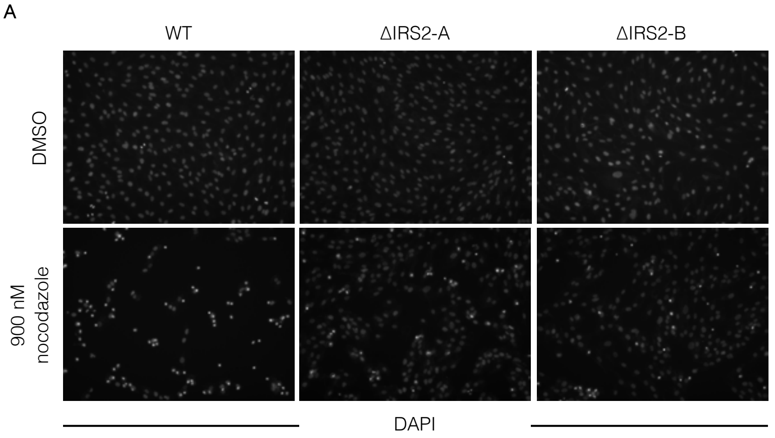


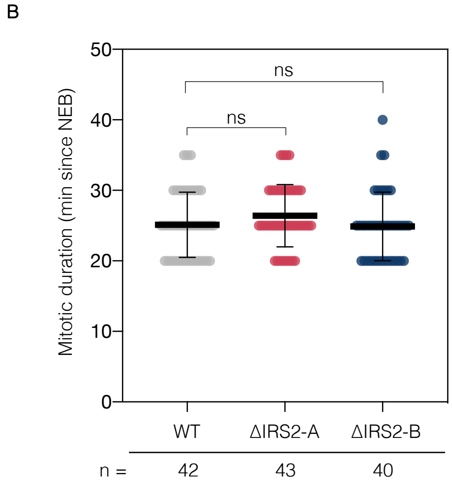


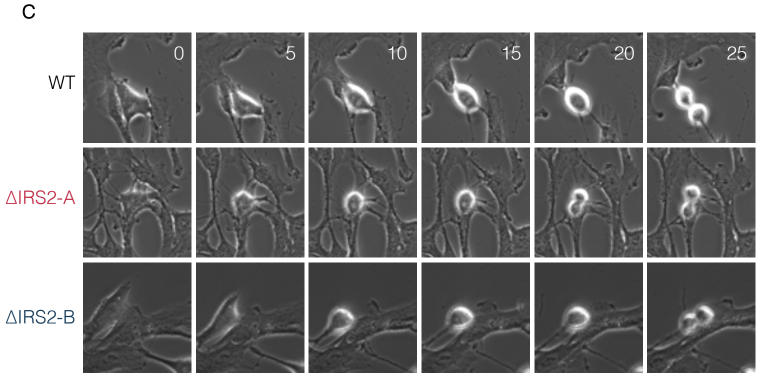

Supplemental Figure 8

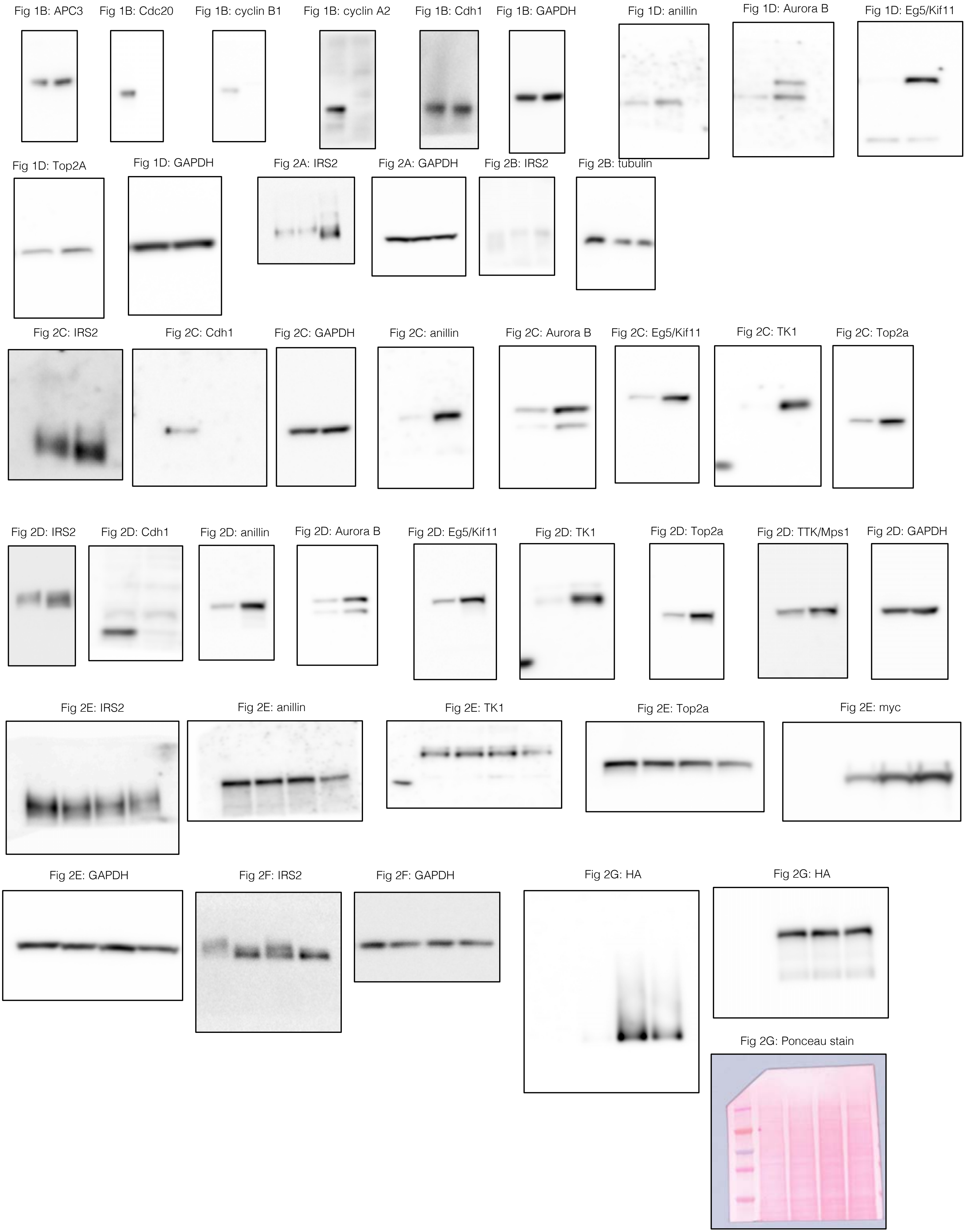

Supplemental Figure 9

Fig 3A: IRS2

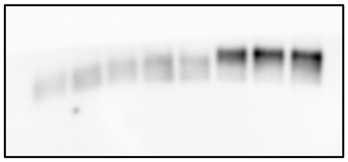

Fig 3: APC3

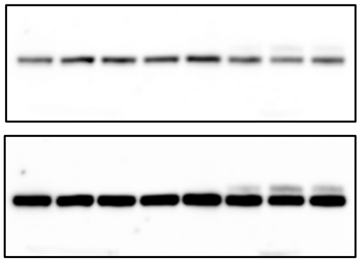

enhanced contrast

Fig 3A: Cyclin B1

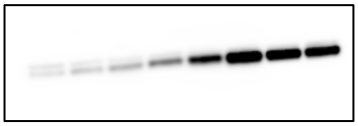

Fig 3A:GAPDH

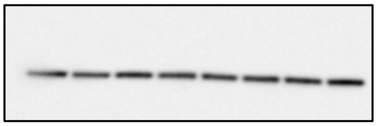

Fig 3B: IRS2

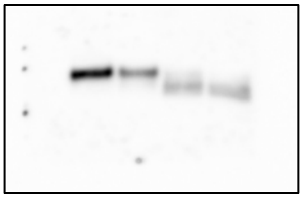

Fig 3B: APC3

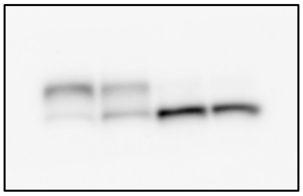

Fig 3B: Cyclin B1

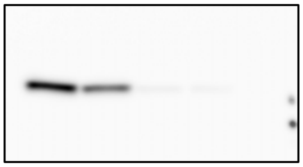

Fig 3B: GAPDH

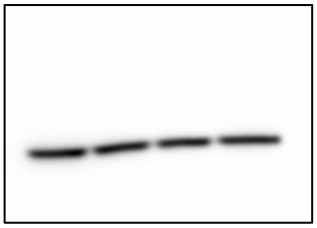

Fig 3C: IRS2

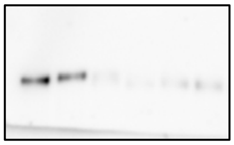

Fig 3C: APC3

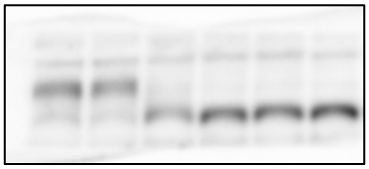

Fig 3C: Cyclin B1

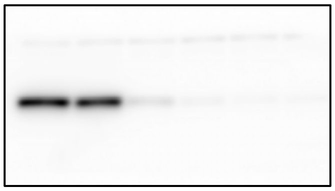

Fig 3C: GAPDH

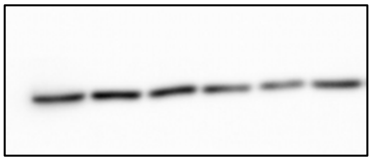

Supplemental Figure 10

Fig 4B: HA

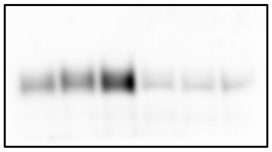

Fig 4B: tubulin

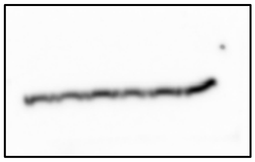

Fig 4C: HA

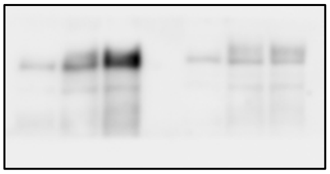

Fig 4C: tubulin

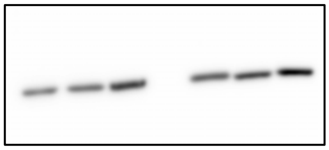

Fig 4D: HA

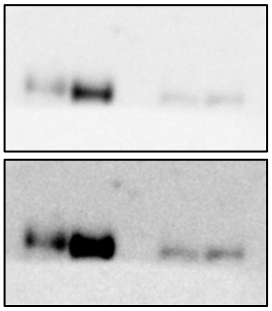

enhanced contrast

Fig 4D: Cdh1

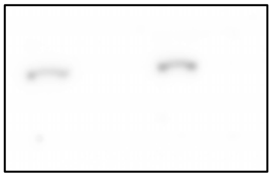

Fig 4D: vinculin

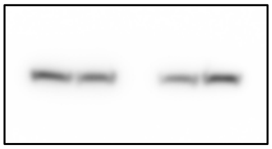

Fig 4E: HA

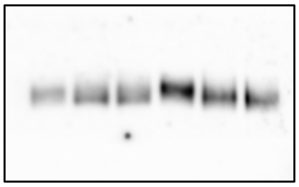

Fig 4H: IRS1

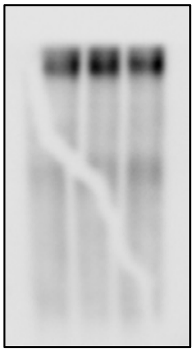

Fig 4H: GAPDH

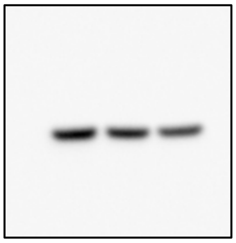

Fig 4E: Cdh1

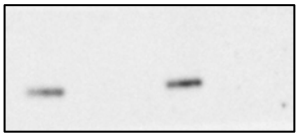

Fig 4E: GAPDH

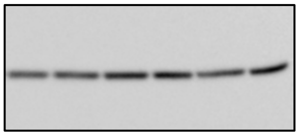

Fig 4I: IRS1

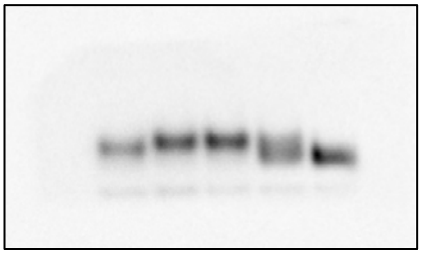

Fig 4I: Cyclin B1

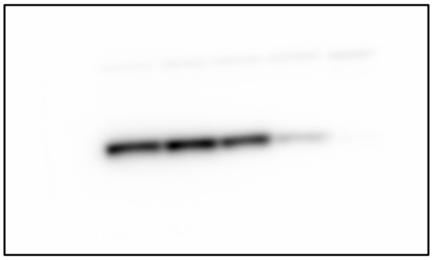

Fig 4I: Cdc20

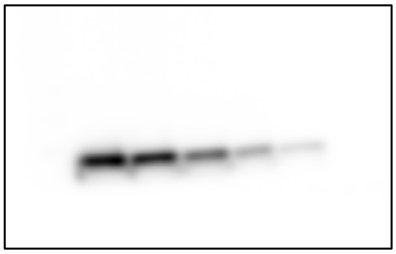

Fig 4I: GAPDH

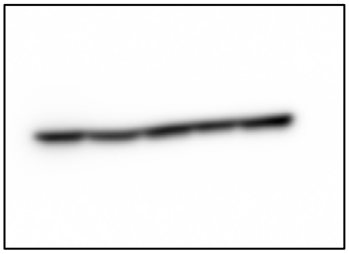

Fig 5A: IRS2

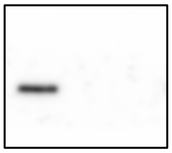

Fig 5A: vinculin

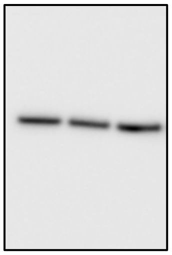
